# MULTIPLE DISTINCT TIMESCALES OF RAPID SENSORY ADAPATION IN THE THALAMOCORTICAL CIRCUIT

**DOI:** 10.1101/2024.06.06.597761

**Authors:** Yi Juin Liew, Elaida D Dimwamwa, Nathaniel C Wright, Yong Zhang, Garrett B Stanley

**Author notes:** **Correspondence:** Professor Garrett B Stanley, Coulter Department of Biomedical Engineering Georgia Institute of Technology & Emory University 313 Ferst Drive, Atlanta GA 30332-0535 USA. Indicates equal contribution (Yi Juin Liew, Elaida D Dimwamwa).

## Abstract

Numerous studies have shown that neuronal representations in sensory pathways are far from static but are instead strongly shaped by the complex properties of the sensory inputs they receive. Adaptation dynamically shapes the neural signaling that underlies our perception of the world yet remains poorly understood. We investigated rapid adaptation across timescales from hundreds of milliseconds to seconds through simultaneous multi-electrode recordings from the ventro-posteromedial nucleus of the thalamus (VPm) and layer 4 of the primary somatosensory cortex (S1) in male and female anesthetized mice in response to controlled, persistent whisker stimulation. Observations in VPm and S1 reveal a degree of adaptation that progresses through the pathway. Signatures of two distinct timescales of rapid adaptation in the firing rates of both thalamic and cortical neuronal populations were revealed, also reflected in the synchrony of the thalamic population and in the thalamocortical synaptic efficacy that was measured in putatively monosynaptically connected thalamocortical pairs. Controlled optogenetic activation of VPm further demonstrated that the longer timescale adaptation observed in S1 is likely inherited from slow decreases in thalamic firing rate and synchrony. Despite the degraded sensory responses, adaptation resulted in a shift in coding strategy that favors theoretical discrimination over detection across the observed timescales of adaptation. Overall, although multiple mechanisms contribute to rapid adaptation at distinct timescales, they support a unifying framework on the role of adaptation in sensory processing.

**Significance Statement:** Although the perceptual effects of persistent sensory stimulation have been known for centuries, the rapid sensory adaptation of the underlying neural signaling to these persistent inputs are not well understood. Here, we present evidence for two distinct timescales of adaptation over several seconds across the thalamocortical circuit in mice. We identify both the overall level of neural activity and the corresponding population synchrony of the thalamic inputs to primary somatosensory cortex as key role players shaping the cortical adaptation.

## INTRODUCTION

The natural sensory environment consists of inputs that span a wide range of spatiotemporal scales. As sensory signals move from the periphery to sensory cortex, sensory representations are progressively transformed via the characteristics of circuitry at each stage and the nature of the anatomical projections across regions (Maravall et al., 2007, 2013; Adibi and Lampl, 2021). Adaptation has been well-documented across all sensory modalities (Whitmire and Stanley, 2016), thought to overcome changes in stimulus statistics via multiple mechanisms that modulate neuronal response properties (Adibi et al., 2013; Latimer et al., 2019). It has long been asserted that the brain’s ability to adapt is a fundamental function that is critical for survival, providing a functional advantage by shifting coding strategies in an ethological advantageous manner (Barlow, 1961). While adaptation has been proposed as a unifying theoretical framework that may integrate numerous disparate observations (Drew and Abbott, 2006), the majority of studies of neuronal adaptation focus on a specific single timescale (Chung et al., 2002; Khatri et al., 2004; Kheradpezhouh et al., 2017). What is not well understood is the evolution of adaptation along different timescales across brain regions in the processing hierarchy, and the functional implications.

In the rodent whisker-mediated somatosensory pathway, previous studies have shown that the degree of sensory adaptation increases as neural signals traverse from periphery to cortex (Ahissar et al., 2000; Khatri et al., 2004; Ganmor et al., 2010; Mohar et al., 2013, 2015). A particularly important stage of the sensory signaling pathway is the thalamocortical circuit, where highly divergent thalamic afferents synapse on to both excitatory and inhibitory neurons in the cortex, initiating thalamic-driven feedforward inhibition that has profound effects on sensory representations (Simons and Carvell, 1989; Gabernet et al., 2005; Cruikshank et al., 2007; Wang et al., 2010a). Thalamocortical synapses exhibit short-term depression, modulating the relative role of the projecting thalamic input (Chung et al., 2002; Castro-Alamancos, 2004a; Katz et al., 2006). However, thalamic neurons themselves also exhibit moderate adaptation, resulting in a withdrawal of the primary drive to cortex. Recent work at the population level also suggests that adaptive changes in thalamic population spiking synchrony play an important role in mediating cortical adaptation through the highly timing-sensitive integration by thalamo-recipient cortical neurons (Wang et al., 2010b; Wright et al., 2021). Given the complexity of the thalamocortical circuit, it is likely that multiple mechanisms are engaged in the transformation of sensory representations throughout the pathway. How this myriad of elements collectively evolves across timescales remains unknown.

Here, we investigated in detail the effects of persistent tactile stimuli on neuronal population activity across the thalamocortical circuit, and specifically the relative roles of aspects of the thalamic input in shaping the cortical response. We conducted multi-site, multi-electrode extracellular recordings of spiking activity from the ventro-posteromedial nucleus of the thalamus (VPm) and layer 4 of the whisker response region of primary somatosensory cortex (S1) in anesthetized mice. We presented controlled, repetitive, punctate whisker deflection trains to individual whiskers and quantified the adaptive effects on single-unit spiking activity in VPm and putative excitatory and inhibitory S1 neurons, as well as in a subset of putative synaptically-connected thalamocortical pairs. In both the cortex and the thalamus, we uncovered evidence of two distinct timescales in the rapid adaptation dynamics over a timecourse of seconds that exhibited differential functional properties across the thalamocortical circuit. We observed an evolution in the relative roles of individual mechanisms across these timescales, with cortical adaptation at the longer timescale appearing to primarily reflect the adaptation of thalamic drive and synchrony. As a result, from the perspective of an ideal observer of spiking activity, adaptation in both of the distinct timescales also continually shifts towards better discrimination of whisker deflection velocity at the expense of detection. These results collectively suggest a complex interplay of adaptive elements that evolve on distinct timescales and position the thalamic population activity as a key role player in shaping cortical representations.

## MATERIALS & METHODS

### Animals

Experiments were conducted using 10 adult C57BL/6J and 14 adult NR133 x Ai32 (NR133 Cre-recombinase driver line crossed with Ai32 mice for expression of channelrhodopsin in the VPm; Bolus et al., 2020; Gerfen et al., 2013) mice of both sexes (14 male, 10 female) aged between 11 and 44 weeks (mean: 20 weeks). All procedures were approved by the Georgia Institute of Technology Institutional Animal Care and Use Committee and followed guidelines established by the National Institutes of Health.

### Head-plate implantation

Head-plate implantation was performed three to seven days before electrophysiological recordings were conducted. A lightweight custom metal (titanium or stainless steel) head-plate was implanted on the skull of the mouse for head fixation and improved stability during recording, in accordance with a previously described protocol (Borden et al., 2017; Liew et al., 2021; Wright et al., 2021; Dimwamwa et al., 2023). During this survival surgical preparation, the animal was sedated with 5% vaporized isoflurane and anesthesia was maintained with 2-3% isoflurane for the remainder of the procedure. An opioid and non-steroidal anti-inflammatory analgesic were administered post-operatively (SR-Buprenorphine 0.8 - 1 mg/kg, SC and Ketoprofen 5-10 mg/kg, IP). The animal’s eyes were covered with ophthalmic eye ointment (Puralube Vet Ointment) to prevent dehydration. Body temperature was monitored and maintained at 37 °C. After sterilizing the skin above the skull and applying topical anesthetic (lidocaine 2%, 0.05 ml, max 7mg/kg, SC), we removed the skin and scraped off periosteum and any conjunctive tissue from the skull. We gently separated tissue and muscles close to the lateral edge of the skull using a scalpel blade (Glass Van & Technocut, size no. 15), leaving sufficient room for head-plate attachment, away from the targeted recording areas. Especially critical for targeting of the ventro-posteromedial nucleus of the thalamus (VPm), the skull surface was leveled during head fixation by adjusting the animal’s head position to minimize the relative height difference (< 100 um) between bregma and lambda (Franklin and Paxinos, 2008). The metal headplate was secured over the intact skull with C&B-Metabond (Parkell Prod Inc) and the rest of the attachment site and edges of the skin incision were secured with skin adhesive (Vetbond, 3M). The final head-plate and dental acrylic structure created a well for the containment of Ringer’s solution (in mM: 135 NaCl, 5 KCl, 5 HEPES, 1 MgCl2-6H2O, 1.8 CaCl2-2H2Ol pH 7.3) for future imaging and electrophysiological recording sessions. We applied a thin layer of transparent Metabond dental acrylic over both hemispheres to prevent inflammation and covered the exposed skull with a silicone elastomer (Kwik-Cast, World Precision Instruments).

### Intrinsic signal optical imaging of the mouse S1

Intrinsic signal optical imaging of whisker-evoked responses in mouse primary somatosensory cortex was performed through the skull under 1-1.2% isoflurane anesthesia to functionally generate a map of the precise location of S1 barrel columns (Lefort et al., 2009; Pala and Petersen, 2015). The imaging was conducted in a separate session to the anesthetized electrophysiology, either within the same session as head-plate implantation or in a separate session. The skull was thinned using a dental drill (0.1 mm diameter bit, Komet, USA) until blood vessels became visible under warm Ringer’s solution. The Ringer’s was then covered with a glass coverslip (thickness 0.13-0.17mm, Fisherbrand). A reference image of the blood vasculature was captured under green illumination (530nm; M530F1 LED, Thorlabs). Repetitive whisker stimuli (4 seconds of stimuli at 10 Hz; sawtooth waveform; 1000 deg/s) were delivered to individual whiskers (see “Sensory stimulus section” below) while imaging S1 under red illumination (625 nm; M625F1 LED, Thorlabs). Images were acquired using a CCD Camera (MiCam02HR, SciMedia) at 100 Hz. Using previously published methods, we recorded intrinsic signals from at least three columns and estimated the location of the unmapped columns by overlaying a template S1 barrel map reconstructed from histology (Liew et al., 2021). At the end of the imaging session, we applied transparent Metabond dental acrylic directly on the S1 site to protect the thinned skull from contamination and infection and sealed the exposed skull with a silicone elastomer (Kwik-Cast, World Precision Instruments).

### Anesthetized electrophysiology

Animals were sedated with 5% vaporized isoflurane and maintained with 2-3% isoflurane. Their eyes were covered ophthalmic eye ointment and the silicone elastomer removed from the skull surface. The skull above the VPm and S1 was carefully thinned with a dental drill (0.1 mm diameter bit, Komet, USA), while frequently irrigating with Ringer’s solution frequently to remove debris and prevent overheating. When the skull was thin enough to easily puncture, we made small craniotomies above VPm and S1. The craniotomies were constantly irrigated with Ringer’s solution to maintain the integrity of the brain tissue.

The animals were then maintained with 1-2% isoflurane for the electrophysiological recordings. Either in separate sessions or in the same session for paired recordings, 32-channel silicon probes with the ‘Poly-3’ configuration (A1x32-Poly3-5mm-25s-177, A1x32-Poly3-6mm-50s-177, or A1x32-Poly3-10mm-50s-177, Neuronexus, Ann Arbor, MI) were independently lowered into the VPm and S1. All signals were acquired using the Tucker Davis Technologies RZ2 Bioprocessor (Alachua, FL). Neuronal signals were amplified, bandpass filtered (500 Hz – 5 kHz), and digitized at 24,414.0625 Hz per channel. Stimulus waveforms and other continuous data were digitized at 10 kHz per channel.

#### VPm recordings

For VPm recordings, the electrode was inserted perpendicular to the cortical surface and typically positioned 1.8 mm caudal and 1.8 mm lateral to bregma and lowered to a depth of 3 – 3.5 mm. If VPm could not be functionally located using this method, we also targeted VPm using coordinates 1.0 - 1.2 mm from the B1 or Beta columns of S1 as identified via the intrinsic signal optical imaging map.

#### S1 recordings

For S1 recordings, a small craniotomy was created at the desired location based on the intrinsic signal optical imaging map, being careful to remain distant from blood vessels. The electrode was positioned at a 35° angle from the vertical axis (parallel to S1) and cortical neurons were recorded at a stereotaxic depth of 300-600 um from the cortical surface, corresponding to layer 4 (L4).

#### Barreloid/Barrel targeting

Any barreloid and/or barrel corresponding to whiskers in the first 4 columns (1-4) and any row (A-E) were considered for recordings, and multiple recordings were conducted per mouse to target multiple barreloids/barrels. First, candidate principal whiskers (PWs), or the whisker that elicits the earliest and largest response for a given barreloid/barrel, were identified through manual stimulation. Then, the responses to various sensory stimuli (see “Sensory stimulus” below) were methodically recorded during computer-controlled galvo-motor stimulation for 3-4 separate whiskers and the principal whisker for the recorded neurons was quantitively determined in post-processing (see “*Principal whisker (PW) identification”* below).

After each recording session, the probes were removed, exposed tissue was covered with agarose, and the well was sealed with Kwik-Cast. Subsequent recording sessions were done every other day. One recording session was conducted per day and two to three recording sessions were conducted for each mouse using either the original craniotomy or additional craniotomies targeting a different cortical column and barreloid.

### Sensory stimulus

Controlled, single whisker stimulation was delivered to the identified whiskers of interest in the rostral-caudal plane using a computer-controlled galvo-motor (galvanometer optical scanner model 6210H, Cambridge Technology). Whiskers were trimmed ∼12 mm from the face. The whisker was fed into the insertion hole at the end of an extension tube (inner diameter: ∼200-300 µm um, length: 15mm) that was connected to the rotor of the galvanometer stimulator. The stimulus probe was positioned 10 mm from the Vibrissal pad. The range of motion of the galvanometer was ± 20°, with a bandwidth of 200 Hz. The galvanometer system was controlled using a custom developed hardware/software system (MATLAB Real-Time Simulink System, MathWorks or the Real-time experiment Interface application (http://rtxi.org/), with a 1 kHz sampling rate. The inter-trial interval (time between end of one trial and beginning of the next) for all stimuli was at least 15 seconds, to minimize any residual effects from the previous trial.

#### Adapting stimulus

To induce rapid sensory adaptation, a train of punctate whisker deflections was presented at 10 Hz (i.e. 100 ms inter-deflection interval) for 10 seconds in the rostral-caudal plane using the galvanometer as described above. Each punctate deflection within the adapting train consisted of a 300 deg/s sawtooth waveform with an exponential rise and decay lasting 17 ms. The reported velocity is defined by the average velocity over the 8.5 ms rising phase (Wang et al., 2010). The neuronal sensory response for control, 1.5 sec adapted, or 10-second adapted was then quantified for the first, 15^th^, or 100^th^ stimulus in the train.

For the assessment of velocity sensitivity as well as the detectability and discriminability from the perspective of an ideal observer, the response to probe stimuli is what is analyzed. Probe stimuli consist of single sawtooth deflections delivered at one of five velocities: 50, 300, 450, 600, 1200 deg/s. The probe deflections were presented with either no preceding adapting stimulus (Control) or with a preceding 1.5 second adapting stimulus (15 deflections at 10 Hz) or 10 second adapting stimulus (100 deflections at 10 Hz). The probe was delivered 100 msec following the end of the adapting trains. All stimulus conditions were randomized across trials, and at least 20 trials per stimulus condition were typically obtained for different velocities.

#### Weak sinusoidal stimulus

To assess monosynaptic connectivity between VPm and S1 neurons, the thalamocortical circuit was probed using a weak, desynchronizing sinusoidal whisker stimulus to elevate baseline firing rates in recorded neurons (Liew et al., 2021). Specifically, 2 - 4 Hz weak sinusoidal deflections of 4 deg amplitude and 4 sec duration (Bruno and Simons, 2002; Bruno and Sakmann, 2006; Wang et al., 2010b) were delivered for approximately 200-500 trials to obtain at least 2000 spikes for the cross-correlation analysis. Occasionally, the first 0.5 - 1 seconds of the trials were eliminated due to high firing rates at the onset of stimulus presentation.

### Optogenetic stimulation of neural activity

A subset of recordings was performed on NR133 x Ai32 transgenic mice to directly activate VPm neurons, bypassing sensory activation from the periphery. In these experiments, a 32-channel silicon opto-electrode with an attached 105 um fiber (A1x32-Poly3-5 mm-25s-177-OA32LP Neuronexus, Ann Arbor, MI) coupled to a 200 μm optic fiber (M95L01, Thorlabs) and a 470 nm LED (M470F3, Thorlabs) was lowered into VPm. The optogenetic stimulation was designed to emulate the adaptation induced by the sensory stimulation described above. The optogenetic adapting stimulus consisted of 470 nm LED pulses that were 17 ms in duration and were delivered to VPm cell bodies for 10 seconds at 10 Hz.

### Post-mortem histology

To verify the recording sites in VPm and S1, a subset of animals was perfused after the paired recording experiments. To label each probe recording track, the probe was slowly retracted along its axis of entry at the conclusion of recording and a few drops of DiI (2.5 mg/mL, in ethanol) were applied on the probe. The electrode was then inserted into the same penetration site along the same axis and the probe was left in the brain for at least 15 minutes. The animal was then euthanized with an overdose of sodium pentobarbital (euthasol, 0.1 mL at 390 mg/mL), a transcardial perfusion was performed, the brain was extracted, and the brain was fixed in 4% paraformaldehyde (PFA, Electron Microscopy Sciences) overnight. The brain was sliced in 100 um coronal sections. The slices were incubated with DAPI (2 mM in PBS, AppliChem) for 15 min before being mounted on slides with a DABCO solution (1,4-Diazabicyclo[2.2.2]octane, Sigma). The slices were imaged with a confocal microscope (Laser Scanning Microscope 900, Zeiss, Germany) to verify the overlap between the DiI stained recording track and the CO-stain.

### Analytical Methods

#### Spike Sorting

Spike sorting was performed offline using KiloSort2.5 (Pachitariu et al., 2024) followed by manual curation of neuronal clusters in Phy2 (Rossant and Harris, 2013). Well-isolated single units were classified using the following criteria: mean waveform signal-to-noise ratio greater than 3 and less than 1% and 3% of all spikes violating the 2 ms refractory period for cortical and VPm neurons, respectively (Liew et al., 2021; Dimwamwa et al., 2023).

#### Analysis of sensory response

For each neuron, only the whisker response to the identified PW was analyzed (see “*Principal whisker (PW) identification*” above). Whisker evoked activity was quantified within the neural response window for each neuron, defined as a 30 ms window following stimulus onset. Evoked response amplitudes were characterized from the trial-to-trial spike count in the neural response window following each deflection. When applicable, evoked responses were normalized to the maximum response to the first three stimuli. The normalized evoked response amplitude was defined as the spike count in the response window divided by the maximum response amplitude evoked by the first three deflections in the adapting trains. The adaptation ratio was computed for each neuron as the average spike count elicited by the last three deflections, normalized to the averaged response to the first three deflections. First-spike latencies were defined as the average time of the first spike within the response window across trials in which there was a spike within the 30 ms response widow (Storchi et al., 2012).

#### VPm neuron validation

It is important to take great care to distinguish different thalamic nuclei from one another, particularly in the mouse where the regions are very small and close together. Thus, all analyses only considered neurons that were responsive to the deflection of at least one whisker. Whisker responsiveness was determined by performing the Wilcoxon signed rank test on the post-stimulus spiking in the 30 ms neuronal response window compared to an equivalent duration pre-stimulus window. Then, a combination of measures was used to classify whisker responsive neurons as originating from the VPm. First, for isolated (non-adapting) stimuli, we quantified the first-spike latency. Second, we quantified the shift in first-spike latency response to adapting stimuli, defined as the time difference between the first-spike latency to the first and last adapting stimuli (Masri et al., 2008). Third, we calculated the response reliability, defined as the percentage of trials where a response was detected within 20 ms in response to each deflection of the repetitive stimulation (8 – 10 Hz; Mainen and Sejnowski 1995). Only neurons with an average first-spike latency of less than 12 ms, latency shift of less than 20 ms, and response reliability of more than 20% were included (Liew et al., 2021; Wright et al., 2021; Borden et al., 2022; Dimwamwa et al., 2023).

#### Cortical neuron classification

S1 layer 4 is the major thalamocortical recipient of VPm projections, where a diversity of cell types receive the feedforward synaptic inputs (Bruno and Simons, 2002). Here, cortical neurons were classified as regular-spiking (RS) or fast-spiking (FS) based on the time interval between the trough and peak of the mean spike waveform, with a cutoff of 0.4 ms separating the broad RS neurons from the narrow FS neurons (Barthó et al., 2004).

#### Principal whisker (PW) identification

Of the 3-4 whiskers chosen for computer-controlled galvo-motor stimulation, the PW was assigned based on the whisker that elicited the largest peak and smallest latency sensory response in the 30 ms neuronal response window. Each VPm neuron was individually assigned a PW using the LFP response for the channel to which the neuron belonged. All simultaneously recorded S1 neurons were collectively assigned a PW based on the MU response across all channels. The sensory responses presented throughout the manuscript include only trials in which each neuron’s primary whisker was being deflected.

#### Determining topographic alignment

A VPm-S1 neuron pair was verified to be topographically aligned when they shared the whisker that elicits the maximum response to isolated stimuli (both spike evoked response and LFP amplitude) and the mean spike latency difference in the sensory response between VPm and S1 neuron pair was between 1-5 ms.

#### Characterization of the adaptation dynamics with exponential models

The adaptive reduction in response amplitudes was characterized by either single or bi-exponential models applied to the normalized evoked response or synchrony over the 10 second adapting stimulus period for the response of either individual neurons or the mean population response. Normalization was to the maximum response/synchrony for the first three stimuli.

For a single exponential fit, the following exponential function was used:

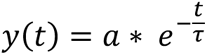

where y(t) is the normalized evoked response at time t, *τ* is the time constant of adaptation, and *a* is a scaling factor.

For the bi-exponential fit, the following function was used:

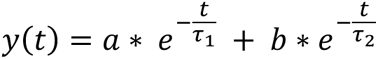

where y(t) is the normalized evoked response at time t, *τ*_1_ and *τ*_2_are the time constants of two timescales of adaptation, and *a* and *b* are scaling factors (Patterson et al., 2013). Importantly, the extra degree of freedom in bi-exponential functions could result in an improvement in model fit regardless of the underlying structure in the data. Thus, we imposed a constraint requiring that the second time constant be at least 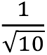 times larger than the first time constant, empirically determined to separate timescales. As a result of this constraint, a pure single exponential would be equally well fit by a single and bi-exponential function.

Note that all whisker responses were analyzed using pure single or bi-exponential models to extract the time constants of the adaptation dynamics, where there was a clear decreasing trend even at the end of the 10 second duration. For the optogenetic responses, however, it was clear that the response plateaued at a non-zero level. Consequently, the analysis for the optogenetic responses focuses on comparing the goodness of fit between the single-exponential model with a constant term and the constrained bi-exponential model. The following function was used to fit the single exponential model with a constant:

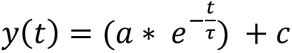

where *c* is the constant.

All parameters were optimized using the ‘fmincon’ function in MATLAB. The optimization aimed to minimize the sum of squared differences between the observed data points and the model predictions. The goodness of fit of each model was quantified by assessing the correlation coefficient (R^2^) between the experimental measures and the fit, and different model complexities were assessed by comparing the R^2^ for the different models across the dataset.

To visualize the exponential decay of the adaptation response, we examined the data within a semi-log plot to identify any exponential components present in the data. Each straight-line segment on the plot corresponds to an exponential component. We then performed linear regression on each segment to determine its slope, which represents the time constant of the exponential component.

#### Analysis of velocity sensitivity

To measure the sensitivity of thalamocortical neurons to sensory stimulus velocity, the whisker response was quantified as a function of increasing deflection velocities: 50, 125, 300, 600, 900, 1200 deg/s. These stimuli were presented either in isolation (Control) or immediately following adapting stimuli (1.5 sec or 10 sec), all randomly interleaved.

#### Spike cross-correlation analysis

Spike cross-correlation analysis was performed to infer monosynaptic connectivity and compute synaptic efficacy for paired recordings in topographically aligned thalamocortical regions, as well as to compute VPm population synchrony. For each spike in a ‘reference’ neuron, a cross-correlogram (CCG) histogram of the relative times of the spikes from a ‘target’ neuron was generated within a 25 msec window before and after each reference spike. All CCGs were constructed using 0.5 ms bins.

#### Inferring monosynaptic connectivity

Using a previously developed method for monosynaptic connectivity inference, in paired and topographically aligned recordings, the thalamocortical circuit was externally driven with a weak whisker stimulus (see Methods: Weak sinusoidal stimulus; Liew et al., 2021). For each topographically aligned VPm-S1 pair, a CCG of the S1 neuron’s spiking relative to the VPm neurons spiking during the weak whisker stimulus was constructed, referred to as the raw CCG. To correct for correlated, stimulus-locked spiking, a shuffle-corrected (SC) spike cross-correlogram was generated. Specifically, a shuffled CCG was generated by randomly shuffling the trials of the VPm and S1 neuron spiking activity relative to each other. The SC CCG was then generated by subtracting the shuffled CCG from the raw CCG. This effectively corrects for any elements of the CCG that are due to the stimulus. To conclude that a neuronal pair was putatively ‘connected’, two criteria were adopted that were expanded from previous studies (Reid and Alonso, 1995; Swadlow and Gusev, 2001; Bruno and Sakmann, 2006) based on both the raw and shuffle-corrected cross-correlogram: (Criterion 1) a notable sharp, millisecond-fine peak observed within a narrow lag of 1 - 4 ms after a VPm spike that is the maximum in the raw CCG; and (Criterion 2) this fast ‘monosynaptic peak’ is significant or still present after accounting for (subtracting) stimulus-induced correlations in the SC CCG (Liew et al., 2021). Because this approach is statistical in nature, and not definitive, we refer to such pairs as ‘putative’ monosynaptically connected pairs.

#### Measuring synaptic efficacy

To quantify the effects of adaptation on synaptic transmission across the thalamocortical junction, we measured synaptic efficacy, quantified as the number of spikes within a 1 - 3 ms window of the SC CCG divided by the total number of VPm spikes used in the analysis (Swadlow and Gusev, 2001). Synaptic efficacy was computed per stimulus in the adapting train using an 80 ms post-stimulus response window.

#### Measuring VPm synchrony

To measure the level of synchronization in the VPm, synchrony was computed for all pairs of simultaneously recorded VPm neurons that responded to the same principal whisker. Synchrony was quantified using the total number of spikes within the central ± 7.5 ms window of the CCG generated from stimulus-evoked spiking in the 30 ms response window divided by the total number of spikes in the raw CCG (Borden et al., 2017; Wright et al., 2021). When applicable, normalized synchrony was normalized to the maximum response to the first three stimuli.

#### Receiver operator characteristic (ROC) analysis

To evaluate the functional consequences of the effects of adaptation over timescales, the theoretical detectability of sensory features was assessed for significantly responsive RS and FS neurons by applying ideal observer analysis (Wang et al., 2010b; Ollerenshaw et al., 2014; Whitmire and Stanley, 2016). The noise distribution was generated from baseline activity, or the spike count 30 ms before the stimulus onset in each trial. For each cortical neuron response to a stimulus at a given velocity, the signal came from a probability distribution estimated using the across-trial mean (μ) and standard deviation (σ) of the spike count per stimulus in the 30 ms neural response window. A parameterized ‘population’ distribution was constructed for each neuron using a Gamma distribution, Γ (N*α, Θ), where α = μ^2^/σ^2^, Θ = σ^2^/μ, and N is the assumed number of identical neurons that share response properties similar to the experimentally recorded neuron. Results were reported using N = 10 but results were qualitatively similar for N = 1, 5, and 20. 1000 samples (trials) were drawn from the ‘population’ distribution.

ROC curves were generated by sliding the threshold across the noise and signal distributions, generating a range of false alarm rates versus hit rates that reflect the degree of overlap of the distributions. The area under the ROC curve (AUROC) was used as a performance metric for theoretical detectability. Note that higher AUROC values indicate better performance as an AUROC value of 1 corresponds to zero overlap between signal and noise distributions (an ideal detector). An AUROC value of 0.5 corresponds to complete overlap between distributions, hence operating at chance.

For theoretical discriminability, the ability of an ideal observer to discriminate between five different velocities (50, 300, 450, 600, 1200 deg/s) was quantified based on the observed S1 activity. S1 recordings of L4 RS and L4 FS neurons with at least 20 stimulus trials per velocity condition were used to generate a pool of 205 L4 RS neurons and 146 L4 FS neurons. Stimulus trials, comprising a minimum of 20 trials per condition, were amalgamated from randomly sampled cortical neurons. These datasets were employed to construct spike count distributions, which were subsequently parameterized using a Gamma distribution. The resulting distributions were input into Linear Discriminant Analysis (LDA) classifiers using MATLAB’s ‘fitcdiscr’ command to evaluate theoretical discrimination performance. This process was iterated 1000 times for each stimulus condition. We conceptualize the ideal observer as an LDA decoder employing maximum likelihood (ML) estimation to allocate an observed response to a specific stimulus velocity. The accuracy of this classification process forms a performance matrix. Given the five stimulus velocities employed here, the performance matrix is a 5×5 matrix where the (i,j) element represents the probability of assigning the observed response to stimulus s_j_ when the actual stimulus is s_i_. The theoretical discrimination performance was quantified as the sum of all diagonal elements of the performance matrix. Given that the five velocities were presented with equal probability, achieving chance discrimination corresponds to a probability of 0.2.

#### Statistical analyses

In order to better visualize the trends in the data, the error bars presented throughout the manuscript represent either the standard error of the mean (SEM) or confidence intervals. However, all comparisons of measures of the neuronal response across stimulus conditions were assessed with the two-sided Wilcoxon signed rank test with a Bonferroni correction to correct for multiple comparisons, which is independent of the visualization. Additionally, comparisons of the decay profile of individual neurons fit with single versus bi-exponential functions were assessed with the one-sided Wilcoxon signed rank test.

### Experimental design

This study consists of a within-subject design such that all comparisons are between the same set of neurons, representative of multiple animals of both sexes (see Animals above).

### Data accessibility

The processed data generated in this study will be made publicly available upon publication.

## RESULTS

### Differential rapid sensory adaptation in VPm and S1

To assess the effects of adaptive sensory inputs on thalamocortical activity, we recorded extracellular neuronal spiking in the vibrissa lemniscal pathway of lightly isoflurane-anesthetized, head-fixed mice using multi-channel silicon probes. We measured the activity of populations of well-isolated, single neurons in the VPm of the thalamus as well as layer 4 of the primary somatosensory “barrel” cortex (S1 L4). S1 neurons were further parsed into putative excitatory regular spiking (RS) and putative inhibitory fast spiking (FS) neurons based on the trough-to-peak time of the spike waveform (Figure 1A; see Methods). We assessed the neuronal responses to a 10 Hz pattern of deflections of a single whisker on the contralateral side of the face using a computer-controlled actuator for up to 10 seconds (each deflection is a transient, 300 deg/s sawtooth whisker deflection; see Methods).

**Figure 1.**
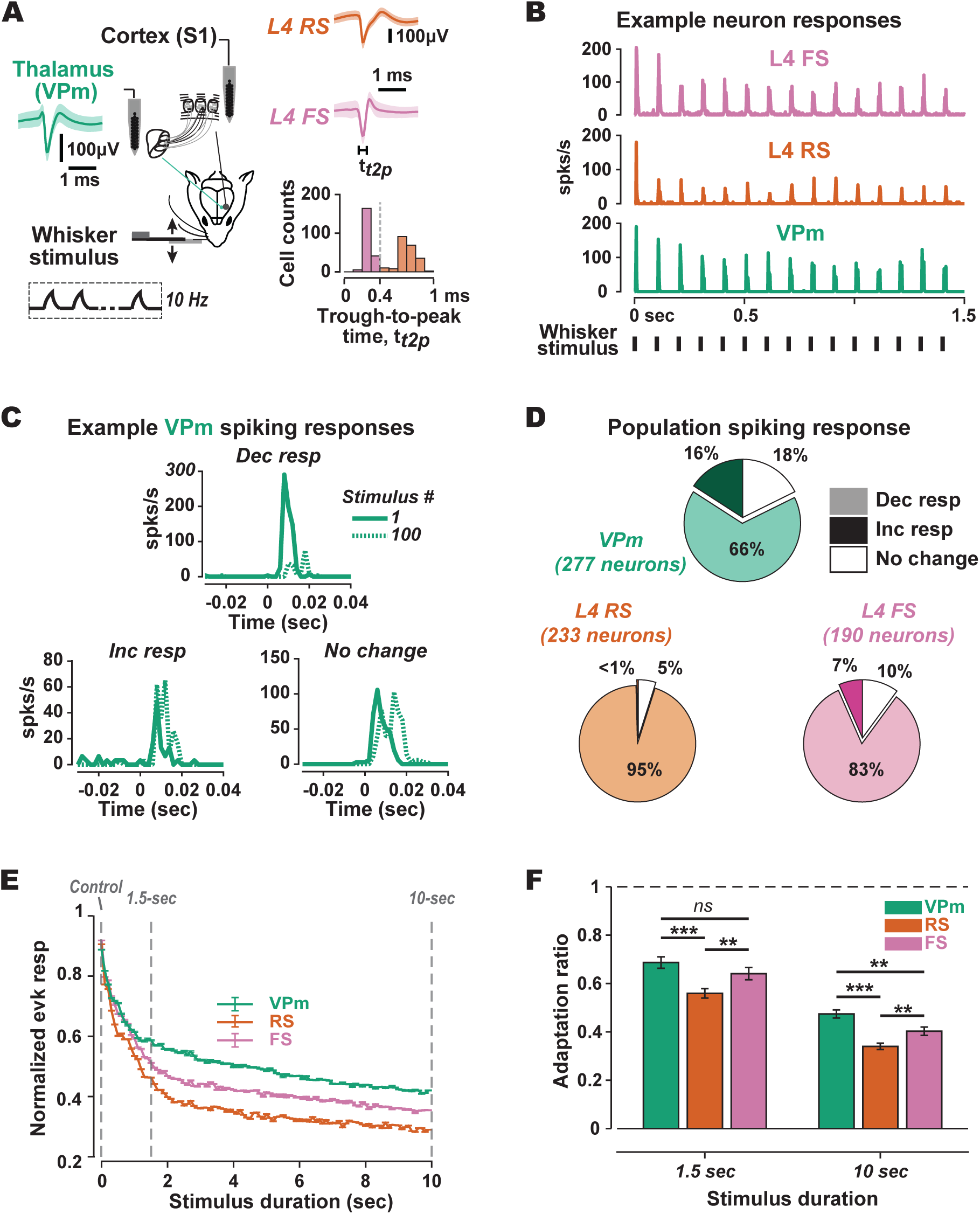
Differential rapid sensory adaptation in VPm and S1. **A**. 32-channel silicon probe recordings were performed in the ventro-posteromedial nucleus of the thalamus (VPm) and in layer 4 (L4) of the primary somatosensory cortex (S1) in lightly isoflurane-anesthetized mice. The mean +/- sem waveforms of example well-isolated VPm, putative excitatory L4 regular spiking (RS), and putative inhibitory L4 fast spiking (FS) typical of each population are shown in green, orange, and pink, respectively. RS and FS cortical neurons were identified based on the trough to peak time of their spike waveforms. Precise deflections were delivered to single whiskers for sensory input using a computer-controlled actuator. Adaptation was induced using a train of sawtooth punctate sensory stimuli (300 deg/s) delivered at 10 Hz for 10 sec (see Methods). **B.** Peri-stimulus time histograms (PSTHs) of the example neurons from A in response to 10 Hz repetitive sensory stimulation. **C.** Overlaid PSTHs of three example VPm neurons’ sensory responses to the 1^st^ and 100^th^ sensory stimulus in the 10 Hz repetitive stimulus train (solid and dashed lines, respectively), each with differing response profiles. The stimulus is delivered at time 0 sec in each plot. **D.** The distribution of responsive neurons from each population that exhibited a decrease, an increase, or no change in sensory response between the 1^st^ and 100^th^ sensory stimulus. Change was determined using the Wilcoxon signed rank test on the per-trials spike count in the 30 ms neural response window for the 1^st^ vs 100^th^ stimulus in the repetitive train. **E.** Grand-averaged mean +/- sem evoked response for the population of VPm, L4 RS, and L4 FS neurons to each stimulus in the 10 second repetitive stimulus train, normalized to the maximum response of the first three stimuli (see Methods). **F.** The mean +/- sem adaptation ratio for each neuron population after 1.5 and 10 sec of repetitive stimulation, defined as the ratio of the mean spike count for the last three stimuli divided by the mean spike count for the first three stimuli. (1.5 sec adapted: VPm: 0.70 ± 0.02, RS: 0.56 ± 0.02; FS: 0.65 ± 0.02; VPm vs RS: p = 2.60e-07, VPm vs FS: p = 6.82e-02, RS vs FS: p = 3.96e-04; 10 sec adapted: VPm: 0.48 ± 0.02, RS: 0.34 ± 0.01; FS: 0.41 ± 0.02; VPm vs RS: p = 1.52e-12, VPm vs FS: p = 8.25e-04, RS vs FS: p = 1.35e-03. All statistical comparisons were performed using the Wilcoxon rank-sum test with a Bonferroni correction; *** indicates p < 3.3e-04, ** indicates p < 3.3e-03, * indicates p < 1.67e-02).

The grand-averaged peri-stimulus time histograms (PSTHs) in Figure 1B show the effects of sensory adaptation induced by 10 Hz repetitive stimulation on the spiking activity of example VPm, S1 L4 RS, and L4 FS neurons. As qualitatively observed for these example neurons, the sensory response of most recorded neurons in each neuronal population decreased over the timecourse of the repetitive stimulation. However, some neurons exhibit no overall change in stimulus-evoked spiking, and others an increase, as demonstrated in VPm by the overlaid PSTHs of the 1^st^ and 100^th^ stimuli in the adapting stimulus train in Figure 1C. All sensory responsive VPm and S1 L4 neurons were therefore parsed into three distinct categories - increased firing, decreased firing, and no change in firing - based on the cumulative spiking evoked within 30 ms of the 1^st^ and 100^th^ deflection of the adapting stimulus. We found that 66%, 95%, and 83% of VPm, RS, and FS neurons, respectively, decreased their sensory response between the 1^st^ and 100^th^ deflection (Figure 1D).

Despite some variability across neurons, we found that most responsive neurons across these regions responded most vigorously to the first few deflections in the stimulus train and adapted strongly to repetitive sensory stimuli. This is evident in the normalized population-averaged evoked response, calculated for each deflection in the train as the trial-averaged number of spikes in the 30 ms response window following the deflection (Figure 1E). The sensory responses were normalized by the un-adapted response, or the maximum response to the first three deflections, hereafter referred to as the control response. For each population, the response amplitude was reduced between 0 and 1.5 seconds, with a more gradual reduction in responsiveness from 1.5 and 10 seconds. Hence, throughout the rest of the study, we exclusively analyzed neurons whose sensory responses decreased between the control and the last stimulus in the 10 second train.

As further evidenced in Figure 1E, L4 RS and FS neurons adapted to a greater degree than VPm neurons in terms of their normalized response amplitude. Consistent with previous studies (Simons and Carvell, 1989; Gabernet et al., 2005; Heiss et al., 2008; Wright et al., 2021), the adaptation was even more pronounced in the RS population compared to both the VPm and the FS population when considering the adaptive decrease relative to the control sensory response. To quantify this, we calculated the adaptation ratio, or the ratio of the mean spike count in response to the last three stimuli in the repetitive stimulus train divided by the mean spike count in response to the first three stimuli. For the first 1.5 seconds of the adapting train, the adaptation ratios were 0.70, 0.56, and 0.65 for the VPm, RS, and FS populations, respectively. At the 10 second time point in the adapting train, the adaptation ratios decreased to 0.48, 0.34, and 0.41 (Figure 1F). Given these observations, we sought to further investigate the adaptation dynamics in both S1 neuron subtypes and VPm thalamus at three time points of interest: the control time point as well as both the 1.5 and 10 second time points.

### Multiple timescales underlie the adaptation dynamics of cortical neurons in response to repetitive sensory stimulation

Figures 2A and 2B show the grand-averaged PSTHs for the population of L4 RS and FS neurons, respectively (note the different vertical scales for RS versus FS). Generally, both cortical sub-populations generated short-latency, transient spiking in response to the control stimulus. There was a qualitatively observable increase in latency and decrease in spiking response as a function of the duration of the adapting stimulus. Indeed, the mean first spike latency for the L4 RS population increased from 12.2 msec to 19.3 msec between the Control and 1.5 second time point, with a further increase to 20.2 msec at the 10 second time point, corresponding to a 58% and 5% increase in latency between the time points. For the L4 FS population, the first spike latency increased from 10.35 msec to 16.51 msec to 18.09 msec, corresponding to a 59% and 10% increase in latency between the time points (Figure 2C). Similarly, the mean evoked response at the three time points of interest decreased from 0.50 to 0.22 to 0.14 spks/stim for the L4 RS population, corresponding to a 56% and 36% decrease in spiking response between the time points and from 0.72 to 0.41 to 0.30 spks/stim for the L4 FS population, corresponding to a 43% and 27% decrease in spiking response between the time points (Figure 2D).

**Figure 2.**
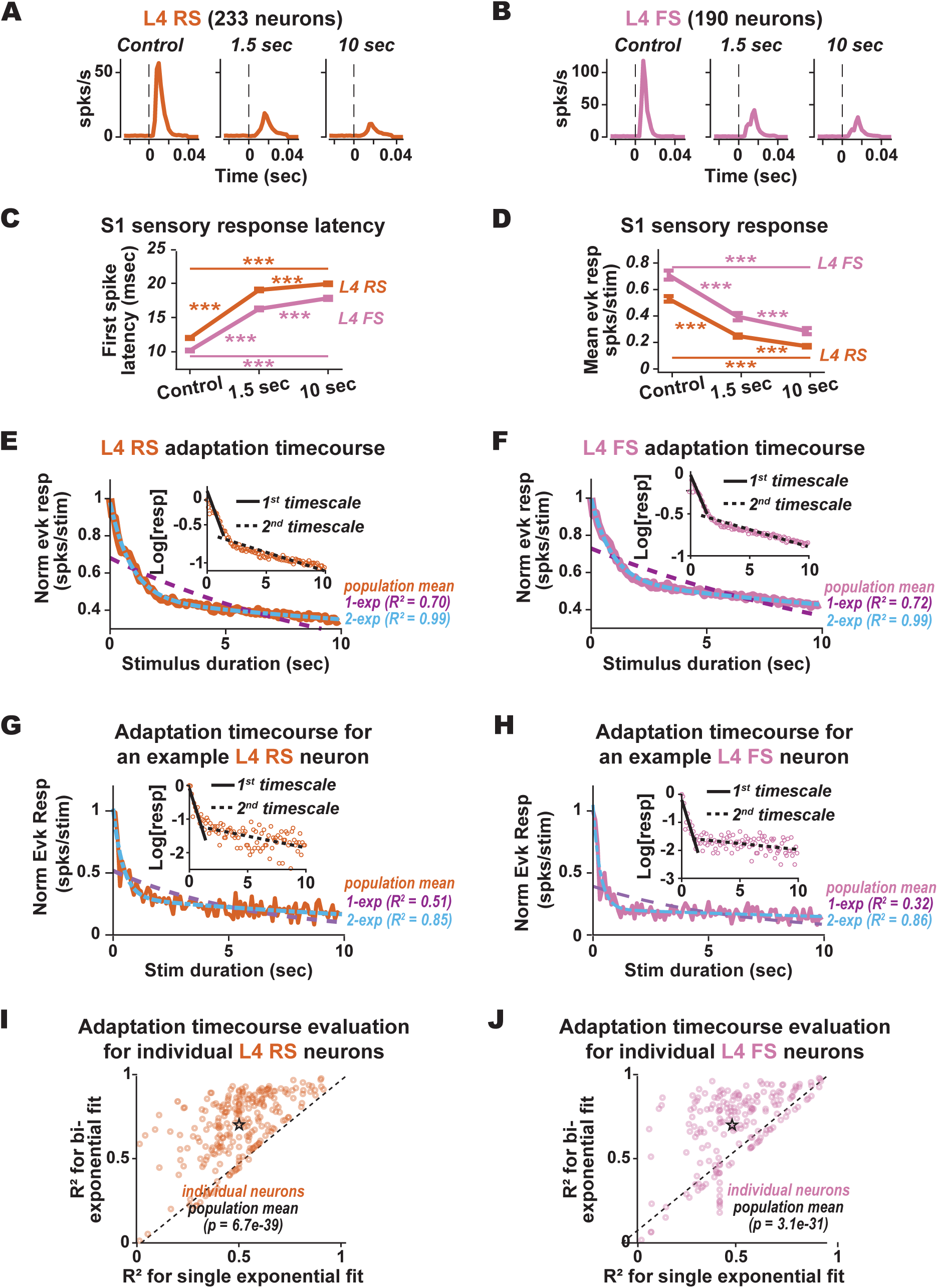
Multiple timescales underlie the adaptation dynamics of cortical neurons in response to repetitive sensory stimulation. **A**. Grand-averaged PSTHs of S1 layer 4 RS neurons to the 10 Hz repetitive stimulus at the three time points of interest: Control (1^st^ stimulus in the repetitive train), 1.5 sec adapted (15^th^ stimulus), and 10 sec adapted (100^th^ stimulus; note that only sensory-responsive neurons whose sensory response significantly decreased between the Control and 10 sec adapted stimuli were included). **B**. Same as A, but for the L4 FS population. **C.** Mean ± sem first-spike latency of the L4 RS and FS populations to repetitive sensory stimuli at the three time points of interest (RS: Control: 12.21 ± 0.15 msec, 1.5 sec adapted: 19.31 ± 0.19 msec, 10 sec adapted: 20.22. ± 0.20 msec; Control versus 1.5 sec adapted: p = 6.40e-40, 1.5 sec adapted versus 10 sec adapted: p = 6.62e-07, Control versus 10 sec adapted: p = 5.85e-40; FS: Control: 10.35 ± 0.13 msec, 1.5 sec adapted: 16.51 ± 0.21 msec, 10 sec adapted: 18.09 ± 0.25 msec; Control versus 1.5 sec adapted: p = 1.04e-31, 1.5 sec adapted versus 10 sec adapted: p = 5.31e-19, Control versus 10 sec adapted: p = 6.26e-33; all statistical comparisons were performed using the Wilcoxon signed-rank test with a Bonferroni correction; *** indicates p < 3.3e-04). **D**. Mean ± sem evoked firing rate to repetitive sensory stimuli at the three time points of interest for the L4 RS and FS populations (RS: Control: 0.50 ± 0.02 spks/stim, 1.5 sec adapted: 0.22 ± 0.01 spks/stim, 10 sec adapted: 0.14 ± 0.01 spks/stim ; Control versus 1.5 sec adapted: p = 1.65e-35, 1.5 sec adapted versus 10 sec adapted: p = 3.36e-31, Control versus 10 sec adapted: p = 7.77e-40; FS: Control: 0.72 ± 0.03 spks/stim, 1.5 sec adapted: 0.41 ± 0.0.03 spks/stim, 10 sec adapted: 0.30 ± 0.02 spks/stim ; Control versus 1.5 sec adapted: p = 7.56e-26, 1.5 sec adapted versus 10 sec adapted: p = 5.48e-28, Control versus 10 sec adapted: p = 7.58e-31; all statistical comparisons were performed using the Wilcoxon signed-rank test with a Bonferroni correction; *** indicates p < 3.3e-04). **E**. The normalized mean evoked response for the L4 RS population over the timecourse of repetitive stimulation overlaid with a single and a bi-exponential model fit (purple and blue dashed lines, respectively; see Methods). The evoked response is normalized per neuron by the maximum response to the first three stimuli before taking the average across all neurons. Inset: semi-log plot of the normalized evoked response. The black lines denote a linear fit over two timescales, suggesting bi-exponential behavior (1_1_ = 0.93 sec, 1_2_ = 37.98 sec). **F**. Same as E, but for the L4 FS population (1_1_ = 0.92 sec, 1_2_ = 40.53 sec). **G**. Same as E, but for an example L4 RS neuron (1_1_ = 0.43 sec, 1_2_ = 26.53 sec). **H**. Same as E, but for an example L4 FS neuron (1_1_ = 0.31 sec, 1_2_ = 29.23 sec). **I**. The correlation coefficients (R^2^) of single vs bi-exponential functions fit to the adaptation timecourse of individual L4 RS neurons (single exponential mean = 0.47, bi-exponential mean = 0.70; the p-value indicates comparison between the single vs bi-exponential model fits assessed for all neurons using the one-sided Wilcoxon signed rank test). **J**. Same as I, but for individual L4 FS neurons (single exponential mean = 0.48, bi-exponential mean = 0.70).

Given the non-saturating nature of the observed adaptation dynamics, we quantified the timescales of the decay in responsiveness of the cortical populations by fitting exponential models to the normalized population-averaged mean evoked response as a function of stimulus duration (see Methods). Although adaptation is widely characterized as a simple, single exponential decay (Chung et al., 2002; Khatri et al., 2004; Patterson et al., 2013; Kheradpezhouh et al., 2017), we found that the declining profile of the adaptation responses over the 10 second repetitive stimulation was better characterized with bi-exponentials (RS: R^2^ = 0.99; FS: R^2^ = 0.99) compared to single exponential functions (RS: R^2^ = 0.70; FS: R^2^ = 0.72; Figures 2E & 2F). This trend was even more evident when plotted on semi-log axes, where the plot reveals two strikingly clear linear regimes with negative slopes, suggesting two distinct timescales of exponential decay (figure insets).

Figures 2G & 2H show the declining profile of the adaptation response of an example L4 RS and L4 FS neurons, respectively. As illustrated by these 2 example neurons, the adaptation timecourse of individual neurons were variable resulting in overall lower R^2^ values, as compared to the fits on the population means. Nonetheless, the responses of individual neurons were also better characterized with bi-exponential functions (RS: R^2^ = 0.85; FS: R^2^ = 0.86) compared to single exponential functions (RS: R^2^ = 0.51; FS: R^2^ = 0.32). Importantly, we used a constrained exponential fitting approach to characterize neuronal adaptation dynamics, ensuring that the second time constant is much larger than the first (see Methods). In cases where the dynamics are purely single exponential, the goodness of fit, R^2^ for the second exponential remains equivalent to that of the first, reflecting the absence of additional slower components. When assessing the exponential fits across all individual L4 RS and L4 FS neurons individually, we find that overall, the sub-populations are significantly better fit by bi-exponential functions than single exponential functions (Figures 2I & 2J), thus directly contributing to two timescales of decay of the population.

### Multiple timescales underlie the adaptation dynamics of thalamic firing rate and population synchrony in response to repetitive sensory stimulation

The clear multi-timescale adaptation dynamics observed in S1 opens the possibility of multiple factors contributing to these observed cortical dynamics. A likely important contributor is the spiking patterns of the VPm neurons that provide the direct input to the cortical layer 4 neurons, in both the mean firing rate and population synchrony of this highly convergent input. Towards characterizing the adaptation dynamics in VPm, Figure 3A shows the grand-averaged PSTHs for the VPm population in response to the adapting sensory stimulus at the three time points of interest. Like S1 neurons, VPm neurons generated short-latency, transient spiking in response to the Control stimulus with shifts in latency and spiking response from the Control to the 1.5 second and 10 second time points. When quantifying these shifts for the population, the mean first spike latency increased from 9.16 to 12.88 to 14.01 msec for the three time points of interest (Figure 3B). These shifts in latency correspond to a 40% increase from Control to the 1.5 second time point, and a further increase of 8.75% between the 1.5 second the 10 second time points. Additionally, the mean evoked response decreased from 0.81 to 0.52 spks/stim between the Control and 1.5 second time point, with an additional decrease to 0.38 spks/stim from the 1.5 second to the 10 second time point (Figure 3C), corresponding to 36% and 27% decreases in sensory response, respectively. Taken together, adaptation dynamics in the VPm also did not saturate between 1.5 to 10 seconds, but rather continued to progress throughout the duration of the adapting stimulus.

**Figure 3.**
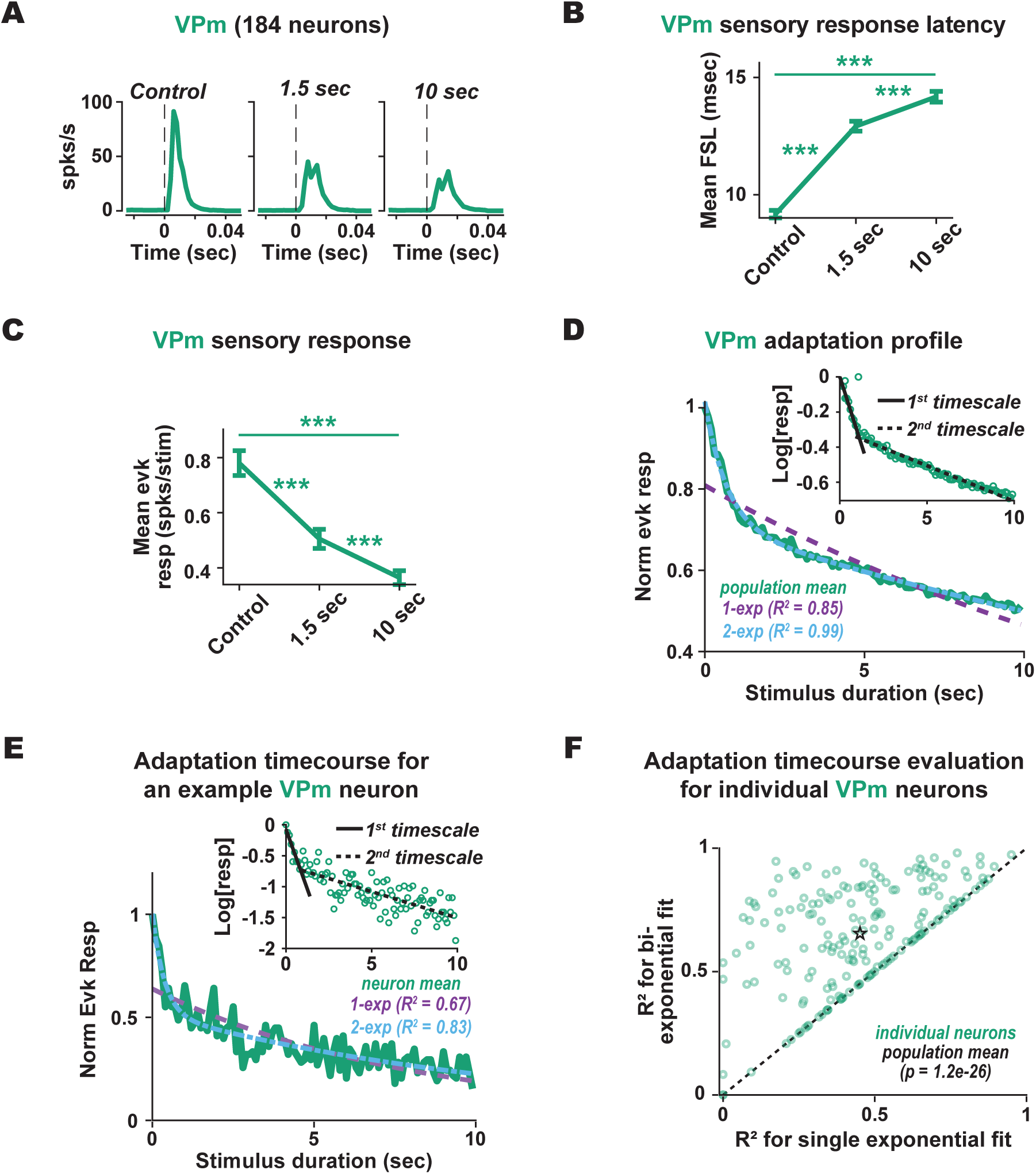
Multiple timescales underlie the adaptation dynamics of thalamic firing rate in response to repetitive sensory stimulation. **A**. Grand-averaged PSTHs of VPm neurons to the 10 Hz repetitive stimulus at the three time points of interest: Control (1^st^ stimulus in the repetitive train), 1.5 sec adapted (15^th^ stimulus), and 10 sec adapted (100^th^ stimulus; note that only responsive neurons whose sensory response significantly decreased between the Control and 10 sec adapted stimuli were included). **B**. Mean ± sem first-spike latency of the VPm population to repetitive sensory stimuli at the three time points of interest (Control: 9.16 ± 0.15 ms, 1.5 sec adapted: 12.88 ± 0.20 ms, 10 sec adapted: 14.01 ± 0.21 ms; Control versus 1.5 sec adapted: p = 1.63e-31, 1.5 sec adapted versus 10 sec adapted: p = 1.39e-17, Control versus 10 sec adapted: p = 1.41e-31; all statistical comparisons were performed using the Wilcoxon signed-rank test with a Bonferroni correction; *** indicates p < 3.3e-04). **C**. Mean ± sem evoked firing rate of the VPm population to repetitive sensory stimuli at the three time points of interest (Control: 0.81 ± 0.05 spks/stim, 1.5 sec adapted: 0.52 ± 0.03 spks/stim, 10 sec adapted: 0.38 ± 0.02 spks/stim. Control versus 1.5 sec adapted: p = 1.15e-13; 1.5 sec adapted versus 10 sec adapted: p = 1.44e-23, Control versus 10 sec adapted: p = 1.43e-27, all statistical comparisons were performed using the Wilcoxon signed-rank test with a Bonferroni correction; *** indicates p < 3.3e-04). **D**. The normalized mean evoked response for the VPm population over the timecourse of repetitive stimulation overlaid with a single and a bi-exponential model fit (purple and blue dashed lines, respectively). The evoked response is normalized per neuron by the maximum response to the first three stimuli before taking the average across all neurons. Inset: semi-log plot of the normalized evoked response. The black lines denote a linear fit over two timescales, suggesting bi-exponential behavior (1_1_ = 0.79 sec, 1_2_ = 29.65 sec). **E**. Same as D, but for an example VPm neuron (1_1_ = 0.37 sec, 1_2_ = 11.96 sec). **F**. The correlation coefficients (R^2^) of single vs bi-exponential models fit to the adaptation timecourse of individual VPm neurons (single exponential mean = 0.45, bi-exponential mean = 0.65; the p-value indicates comparison between the single vs bi-exponential model fits assessed for all neurons using the one-sided Wilcoxon signed rank test).

To characterize the timescale of the decay in responsiveness of VPm neurons, we again fit with exponential models the normalized population-averaged mean evoked response as a function of stimulus duration. We found that the declining profile of the adaptation response over the 10 second repetitive stimulation was better characterized with a bi-exponential (R^2^ = 0.99) compared to a single-exponential function (R^2^ = 0.85; Figure 3D). Again, the bi-exponential nature of the decay became strikingly clear when plotting the normalized evoked responses on semi-log axes, highlighting two linear regions (figure insets).

We next assessed the adaptation timecourse of individual VPm neurons to determine how they contribute to the bi-exponential decay of the population. Figure 3E shows the declining profile of the adaption response of an individual VPm neuron which is also better characterized with a bi-exponential function (R^2^ = 0.83) compared to a single exponential function (R^2^ = 0.67). The adaptation timecourse of individual VPm neurons is overall more variable which results in lower R^2^ values. Overall, however, individual VPm neurons are themselves significantly better fit with bi-exponential functions than single exponential functions (Figure 3F), thus directly contributing to the two timescales of decay observed at the population level.

A number of studies have shown that S1 can be highly sensitive to changes in the synchrony of the convergent thalamic inputs (Pinto et al., 2000; Swadlow and Gusev, 2001; Bruno and Sakmann, 2006; Wang et al., 2010b, 2010a; Ollerenshaw et al., 2014). We thus sought to also characterize the adaptation profile of VPm synchrony. We quantified the synchrony between every pair of simultaneously recorded neurons having maximal responsiveness to the same whisker (i.e. likely in the same thalamic barreloid). Synchrony was computed by generating a cross-correlogram (CCG) and measuring the occurrence of coincident spikes within a +/- 7.5 ms window, normalized by the number of spikes used in the analysis (see Methods, Figure 4A). This normalization accounts for the overall firing rate which is important given that synchrony and firing rate naturally covary. First examining changes in synchrony in response to the adapting stimuli at the three time points of interest, the VPm population synchrony decreased from 0.89 to 0.81 to 0.76 (unitless quantity, see Methods, Figure 4B). Further, as with the mean firing rate, the synchrony evolved in a bi-exponential manner (R^2^ = 0.92) and was not as well fit with a single exponential (R^2^ = 0.75; Figure 4C).

**Figure 4.**
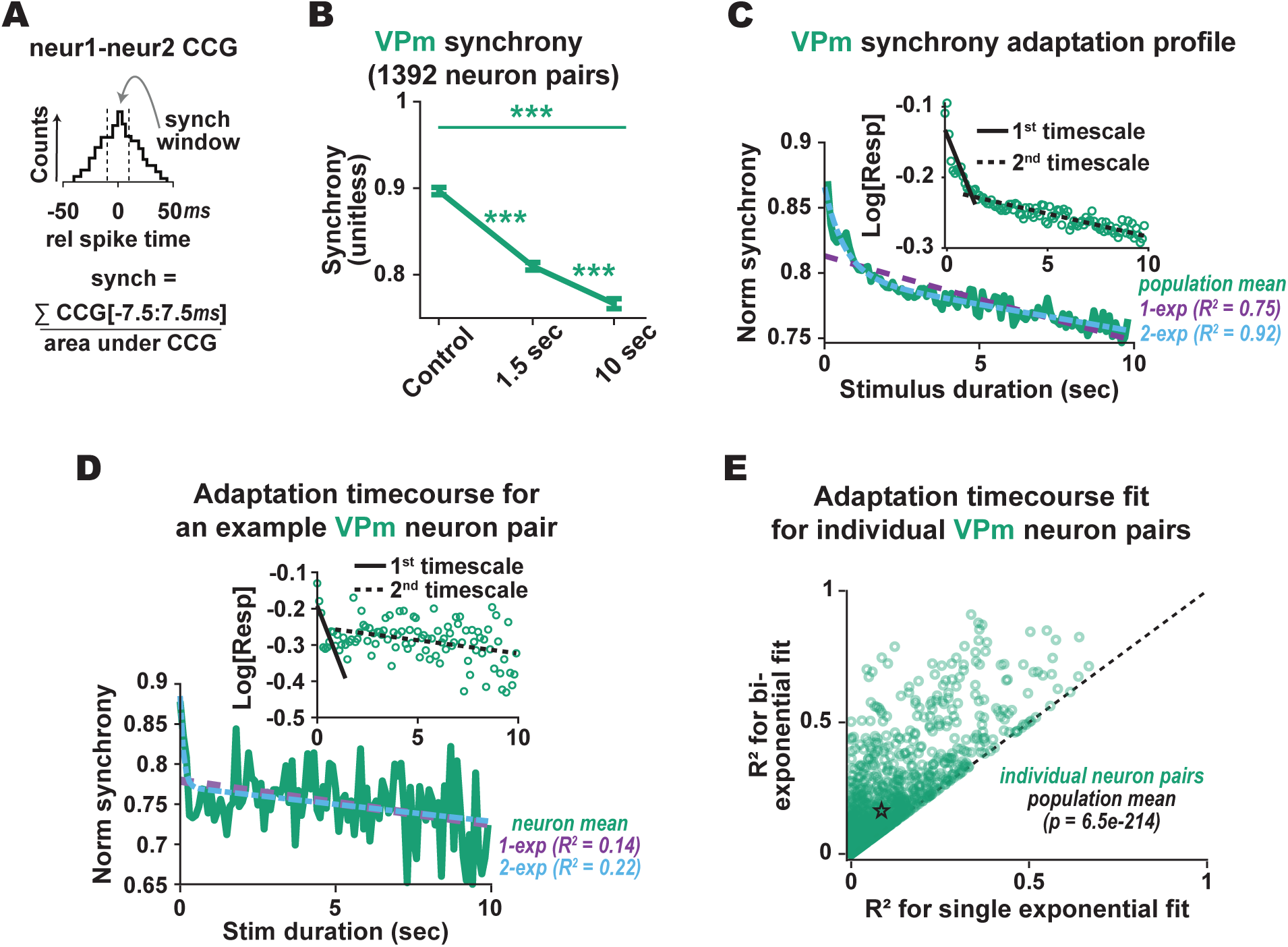
Multiple timescales underlie the adaptation dynamics of thalamic synchrony in response to repetitive sensory stimulation. **A**. VPm population synchrony is measured by generating pairwise cross-correlograms (CCGs) and quantifying the area in the +/-7.5 msec window normalized by the area under the entire CCG (see Methods). **B.** Mean ± sem synchrony of the VPm population to repetitive stimuli at the three time points of interest (Control: 0.89 ± 4.01e-03, 1.5 sec adapted: 0.81 ± 4.30e-03, 10 sec adapted: 0.76 ± 5.52e-03. Control versus 1.5 sec adapted: p = 6.90e-92; 1.5 sec adapted versus 10 sec adapted: p = 1.52e-20, Control versus 10 sec adapted: p = 3.90e-107, all statistical comparisons were performed using the Wilcoxon signed-rank test with a Bonferroni correction; *** indicates p < 3.3e-04). **C.** The normalized mean evoked synchrony for the VPm population over the timecourse of repetitive stimulation overlaid with a single and a bi-exponential model fit (purple and blue dashed lines, respectively). The synchrony is normalized per pair of neurons by the maximum synchrony in response to the first three stimuli before taking the average across all pairs of neurons. Inset: semi-log plot of the normalized population synchrony. The black lines denote the linear fit over two timescales, suggesting bi-exponential behavior (1_1_ = 0.66 sec, 1_2_ = 185.55 sec). **D**. Same as C but for an example pair of VPm neurons (1_1_ = 0.22 sec, 1_2_ = 173.37 sec). **E**. The correlation coefficients (R^2^) of single vs bi-exponential models fit to the adaptation timecourse of individual pairs of VPm neurons (single exponential mean = 0.09, bi-exponential mean = 0.16; the p-value indicates comparison between the single vs bi-exponential model fits assessed for all neurons using the one-sided Wilcoxon signed rank test).

As evidenced in Figure 4D, the adaptation timecourse for the synchrony of individual VPm neuron pairs was also better characterized by bi-exponential functions (R^2^ = 0.22) compared to single exponential functions (R^2^ = 0.14). This example neuron highlights that the adaptation timecourse for the synchrony of individual neuron pairs are variable, owing in part to the data requirements and sensitive nature of synchrony measurements. This in turn results in overall lower R^2^ values for individual neuron pair measures of VPm synchrony than that for measurements of the larger population dataset in Figure 4C. Nonetheless, the synchrony of individual VPm neuron pairs is overall significantly better fit by bi-exponential functions than by single exponential functions (Figure 4E).

### Multiple timescales underlie the adaptation dynamics of thalamocortical synaptic efficacy in response to repetitive stimulation

The above analyses suggest a relationship between cortical L4 neurons and the VPm neurons from which the L4 neurons receive direct sensory input, but do not measure this relationship explicitly. To evaluate the dynamics of the transformation across the thalamocortical synapse, we next examined the effects of the adapting stimulus on synaptic efficacy of putatively connected S1 pairs recorded simultaneously using previously established methods for identifying monosynaptic connectivity in-vivo (Liew et al., 2021). The left panel of Figure 5A shows raster plots of example topographically aligned VPm, L4 RS, and L4 FS neurons that were simultaneously recorded in response to a weak, sinusoidal input stimulus. Note that the overall firing rate of all the neurons was elevated but not generally stimulus-locked; this is critical for identifying putatively connected pairs. The spiking evoked by this weak sinusoidal stimulus was used to generate raw CCGs between every possible pair of topographically aligned VPm-L4 RS neurons. Shuffled CCGs were generated by randomizing the trials of the VPM and S1 spiking activity relative to each other, effectively correcting for any stimulus-induced elements of the CCG. Finally, shuffle-corrected (SC) CCGs were generated by subtracting the shuffled CCGs from the raw CCGs (see Methods). As is the case for the SC CCGs of the example pairs of neurons shown in the right panel of Figure 5A, a narrow peak within the 1-3 msec window with prominence greater than 3.5 standard deviations of the shuffled CCG suggests monosynaptic connectivity (see Methods; Liew et al., 2021). Of the 1199 recorded VPm-RS thalamocortical pairs that were found to be topographically aligned, 297 were inferred to be monosynaptically connected (24.8%). Similarly, of the 681 recorded VPm-FS thalamocortical pairs that were found to be topographically aligned, 319 were inferred to be monosynaptically connected (46.8%). Subsequent analyses were restricted to these putatively connected thalamocortical neuron pairs.

**Figure 5.**
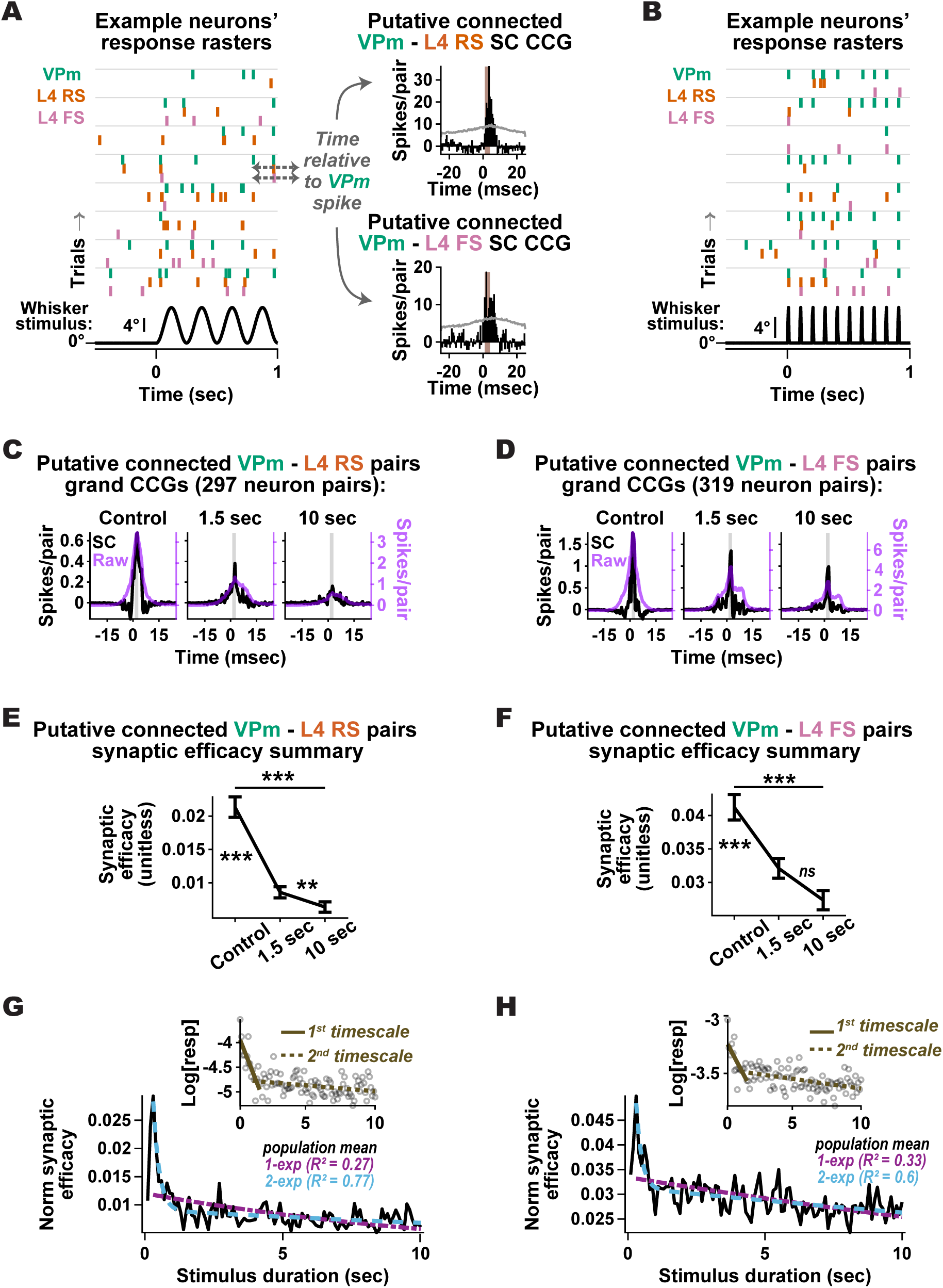
Multiple timescales underlie the adaptation dynamics of thalamocortical synaptic efficacy in response to repetitive sensory stimulation. **A**. Left: Spike raster plots of a simultaneously recorded VPm, L4 RS, and L4 FS neuron in response to a weak sinusoidal whisker stimulus (4 Hz, mean velocity: 25-50 deg/s) in order to elevate spiking activity without driving correlated, stimulus-locked spiking across neurons. Right: The shuffle-corrected (SC) cross-correlograms (CCGs) for the L4 RS (top) and FS (bottom) neuron, each relative to the VPm neuron’s spiking. The SC CCGs are generated by subtracting the shuffled CCGs from the raw CCGs. Raw CCGs are generated using the spikes elicited by the sinusoidal whisker stimulus and the shuffled CCGs are generated by randomizing the trials of the VPM and S1 spiking activity relative to each other. The gray lines denote 3.5 standard deviations of the shuffled distribution. A peak in the SC CCGs in the 1-3 ms window (window highlighted in light brown) indicates that these pairs of neurons are putative monosynaptically connected (Liew et al. 2021). **B**. Spike rasters of the putative monosynaptically connected neurons from A in response to 10 Hz repetitive sensory stimulus (300 deg/s). **C**. The grand-averaged Control (left), 1.5 sec adapted (middle), and 10 sec adapted (right) SC (black) and raw (purple) CCGs for all putative monosynaptically connected VPm-L4 RS neuron pairs. The 1-4 ms window used in subsequent analysis is highlighted in light brown. **D**. Same as C, but for all putative monosynaptically connected VPm-L4 FS pairs. **E**. Mean ± sem synaptic efficacy for all putative monosynaptically connected VPm-L4 RS pairs to repetitive sensory stimuli at the three time points of interest. Given the noisy nature of synaptic efficacy measurements, the control datapoint represents the average efficacy across first the stimuli, the 1.5-sec adapted datapoint represents the average efficacy across the 14^th^-16^th^ stimuli (1.4-1.6 sec), and the 10-sec adapted timepoint represents the average efficacy across the 98^th^-100^th^ stimuli (9.8-10 sec; Control: 2.1e-02 ± 1.7e-03, 1.5 sec adapted: 8.6e-03 ± 9.4e-04, 10 sec adapted: 6.4e-03 ± 8.1e-04. Control versus 1.5 sec adapted: p = 1.2e-11; 1.5 sec adapted versus 10 sec adapted: p = 1.2e-03, Control versus 10 sec adapted: p = 2.2e-15, all comparisons were performed using the Wilcoxon signed-rank test with a Bonferroni correction; *** indicates p < 3.3e-04, ** indicates p < 3.3e-03). Synaptic efficacy is defined for each adaptation time point as the number of synchronous spikes within the 1-3 ms window divided by the number of VPm spikes contributing to the analysis (see Methods). **F**. Same as E, but for all putative monosynaptically connected VPm-L4 FS pairs (Control: 4.1e-02 ± 2.2e-03, 1.5 sec adapted: 3.2e-02 ± 1.9e-03, 10 sec adapted: 2.7e-02 ± 1.9e-03. Control versus 1.5 sec adapted: p = 4.3e-05; 1.5 sec adapted versus 10 sec adapted: p = 3.8e-02, Control versus 10 sec adapted: p = 1.6e-08, all comparisons were performed using the Wilcoxon signed-rank test with a Bonferroni correction; *** indicates p < 3.3e-04). **G.** The normalized mean synaptic efficacy for all putative monosynaptically connected VPm-L4 RS pairs over the timecourse of repetitive sensory stimulation overlaid with a single and a bi-exponential model fit (blue and purple dashed lines, respectively). The efficacy is normalized per neuron by the maximum response to the first three stimuli before taking the average across all neurons. Inset: semi-log plot of the normalized population synaptic efficacy. The purple and blue lines denote the linear fit over two timescales, suggesting bi-exponential behavior (1_1_ = 0.22 sec, 1_2_ = 34.1 sec). **H** Same as G, but for all putative monosynaptically connected VPm-L4 FS pairs (1_1_ = 0.20 sec, 1_2_ = 56.2 sec).

Responses to the 10 Hz adapting stimulus for the same example VPm, L4 RS, and L4 FS neurons in Figure 5A are shown in Figure 5B. For each of the three time points of interest, we assessed the synaptic efficacy of each putatively connected thalamocortical pair, defined as the probability of an S1 neuron spike following a spike of the pre-synaptic VPm neuron. More specifically, using the spiking activity in a 80 msec window following each of the time points of interest in the repetitive stimulus, synaptic efficacy was calculated as the number of synchronous spikes in the SC CCG within the 1-3 ms window following each VPm spike divided by the total number of spikes in the SC CCG (Swadlow and Gusev, 2001).

Figures 5C and 5D show a clear drop in the peak of the grand-averaged SC CCG generated from the spiking responses at the three time points of interest in the 10 Hz repetitive stimulation for VPm-L4 RS and VPm-L4 FS putatively connected pairs, respectively. Indeed, synaptic efficacy decreased across the three time points for both VPm-L4 RS and VPm-L4 FS, summarized in Figures 5E and 5F. Note that the apparent decrease in synaptic efficacy from 1.5 to 10 seconds for the VPm-L4 FS pairs was not statistically significant. When plotting the synaptic efficacy over the timecourse of stimulation, we observe a decaying trend, with an early rapid drop of 81% and 37% at the 1.5 second time point compared to control for the VPm-L4 RS and VPm-L4 FS pairs, respectively. Following this early drop, synaptic efficacy gradually declined during the latter part of stimulation, corresponding to a 50% and 5% drop at the 10 second time point compared to the 1.5 second time point for the VPm-L4 RS and VPm-L4 FS pairs, respectively. Here again, we observe two distinct timescales, as evident in the semi-log plot insets (Figures 5G and 5H). It is important to note that since the above measures of synaptic efficacy were obtained during sensory inputs that synchronize the thalamic population, the efficacy measures could be influenced by the overall drive and synchronization of the thalamic input, as well as depression at the thalamocortical synapse (see Discussion).

### Direct optogenetic drive of VPm thalamus eliminates longer timescale adaptation in VPm

In our next set of experiments, we sought to explore the relationship across the thalamocortical junction through a more direct manipulation of VPm itself. Specifically, we optogenetically stimulated VPm neurons in NR133 x Ai32 mice which enables selective expression of Channelrhodopsin (ChR2) in VPm neurons (Figure 6A). Using an optrode lowered into VPm, we presented pulsatile optogenetic stimulation of identical frequency and duration to the sensory stimulation: 10 Hz for 10 seconds. The LED pulse waveform was designed to elicit a response that mimics the control VPm sensory response (Figure 6B; see Methods).

**Figure 6.**
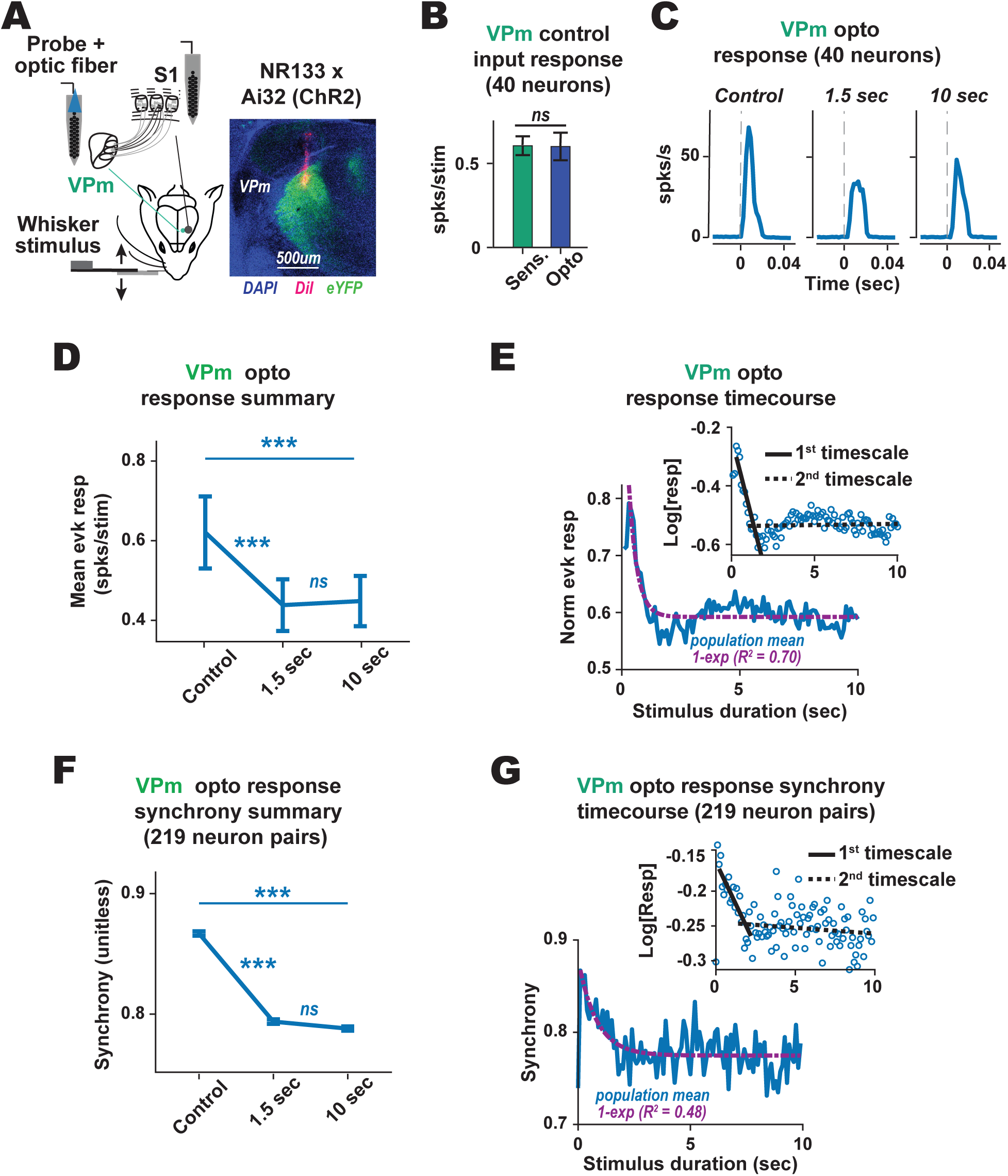
Direct optogenetic drive of VPm thalamus eliminates longer timescale adaptation in VPm. **A**. ChR2-expressing VPm neurons in lightly isoflurane-anesthetized NR133xAi32 mice were activated with 10 Hz repetitive LED pulses to selectively control properties of the VPm inputs while recording VPm and cortical L4 activity using 32-channel silicon probe. The histological section shows expression of ChR2 tagged with eYFP (green) in the VPm and the co-localized electrode track fluorescently labeled with DiI (red) amongst DAPI labelled neurons (blue). **B.** The mean ± sem firing rate of the VPm population to control sensory and opto stimulation (Sensory: 0.6 ± 5.6e-2 spks/stim, Opto: 0.6 ± 0.8.2e-2 spks/stim, p = 0.37, the comparison was performed using the Wilcoxon signed-rank test). **C**. Grand-averaged PSTHs for VPm neurons to 10 Hz opto stimulation of VPm at the three time points of interest. **D.** Mean ± sem evoked response of the VPm population to repetitive optogenetic stimuli at the three time points of interest (Control: 0.62 ± 0.09 spks/stim, 1.5 sec adapted: 0.44 ± 0.06 spks/stim, 10 sec adapted: 0.45 ± 0.06 spks/stim. Control versus 1.5 sec adapted: p = 1.72e-04; 1.5 sec adapted versus 10 sec adapted: p = 7.17e-01, Control versus 10 sec adapted: p = 2.50e-04, all statistical comparisons were performed using the Wilcoxon signed-rank test with a Bonferroni correction; *** indicates p < 3.3e-04). **E.** Population-averaged evoked response for the VPm population over the timecourse of repetitive optogenetic stimulation overlaid with a single exponential model fit. Inset: semi-log plot of the normalized evoked response; the black lines denote the linear fit over one timescale plus a constant, suggesting single exponential behavior (1_1_ = 0.36 sec, constant = 0.61). **F.** Mean ± sem synchrony of the VPm population to repetitive optogenetic stimuli at the three time points of interest (Control: 0.87 ± 1.80e-03, 1.5 sec adapted: 0.79 ± 1.88e-03, 10 sec adapted: 0.79 ± 1.76e-03. Control versus 1.5 sec adapted: p = 1.45e-07; 1.5 sec adapted versus 10 sec adapted: p = 3.48e-01, Control versus 10 sec adapted: p = 8.08e-08, all statistical comparisons were performed using the Wilcoxon signed-rank test with a Bonferroni correction; *** indicates p < 3.3e-04). **G.** Population-averaged synchrony of the VPm population over the timecourse of repetitive optogenetic stimulation overlaid with a single exponential model fit. Inset: semi-log plot of the normalized evoked response; the black lines denote the linear fit over one timescale plus a constant, suggesting single exponential behavior (1_1_ = 0.82 sec, constant = 0.77).

As shown by the grand-averaged PSTHs in Figure 6C, repetitive optogenetic stimulation produced a decrease in response in VPm neurons between the control and 1.5 second time point in the repetitive optogenetic stimulation. Although the optogenetic input induced an initial facilitation in the thalamic response not generally observed in the sensory adaptation, likely due to synchronization effects of the optogenetic light pulse, beyond the first 200-300 ms, the adaptation induced by the optogenetic train was similar to that observed for adaptation with the sensory stimulus (Figure 3A). However, in contrast to the sensory adaptation, qualitatively there was a less clear difference in response between the 1.5 and 10 second time points. In response to the optogenetic stimulation at the three time points of interest, the evoked VPm population firing rate decreased from the Control to the 1.5 second condition but did not further decrease between the 1.5 second and 10 second conditions (Figure 6D). We characterized the timescale of the decay in responsiveness of VPm neurons by again fitting with exponential models the normalized population-averaged mean evoked optogenetic response as a function of stimulus duration. This time, we found that the declining profile was better characterized with a single exponential (R^2^ = 0.70). Naturally, due to the added degree of freedom, the R^2^ value for a bi-exponential is larger (R^2^ = 0.71). However, this increase in R^2^ is marginal and negligible. The single exponential nature of the decay becomes strikingly clear when plotting the normalized evoked responses on semi-log axes, highlighting only one linear region with a slope less than 0, suggesting a single exponential with a constant offset (Figure 6E). It is important to note that the initial quick transient facilitation in the VPm response that was unique to the optogenetic input did not influence this quantification.

Similarly, in response to repetitive optogenetic stimulation, VPm synchrony decreased from Control to the 1.5 second adapted conditions (also similar to that for sensory adaptation – Figure 4B), but no differences were detected between the 1.5 second and 10 second adapted conditions (Figure 6F). In addition, the timecourse of VPm synchrony to repetitive optogenetic stimulation was well fit with a single exponential and not improved with a second exponential, further evidenced by the semi-log insets (Figure 6G; 1-exp: R^2^ = 0.48; 2-exp: R^2^ = 0.48).

### Optogenetic drive of VPm results in a loss of longer timescale adaptation in S1

Having designed a condition that eliminates the longer timescale adaptation in thalamus such that VPm synchrony and firing rate rapidly decayed before stabilizing for the remainder of the repetitive stimulus, we then considered the dynamics in S1. Figures 7A and 7B show the grand-averaged cortical responses to optogenetic stimulation of VPm. The L4 RS and FS neurons adapted strongly to the repetitive optogenetic stimuli. At the three time points of interest, both the L4 RS and FS neuron populations’ firing rate decreased from the Control to the 1.5 second time point but did not further decrease from the 1.5 second to 10 second time point (Figures 7C and 7D, respectively). As evident in the normalized population-averaged mean evoked response in both cortical populations, the reduction in cortical response over the 10 second timecourse was well characterized by a single timescale, with marginal gain for a second exponential (Figure 7E and 7F; 1-exp: R^2^ = 0.78 for L4 RS and R^2^ = 0.81 for L4 FS; 2-exp: R^2^ = 0.78 for L4 RS and R^2^ = 0.83 for L4 FS). When visualizing the timecourse on semi-log axes for both L4 RS and FS populations, a distinct pattern emerges: a single linear region is followed by a relatively flat line. This observation indicates that the decay in evoked responses occurred within a single timescale.

**Figure 7.**
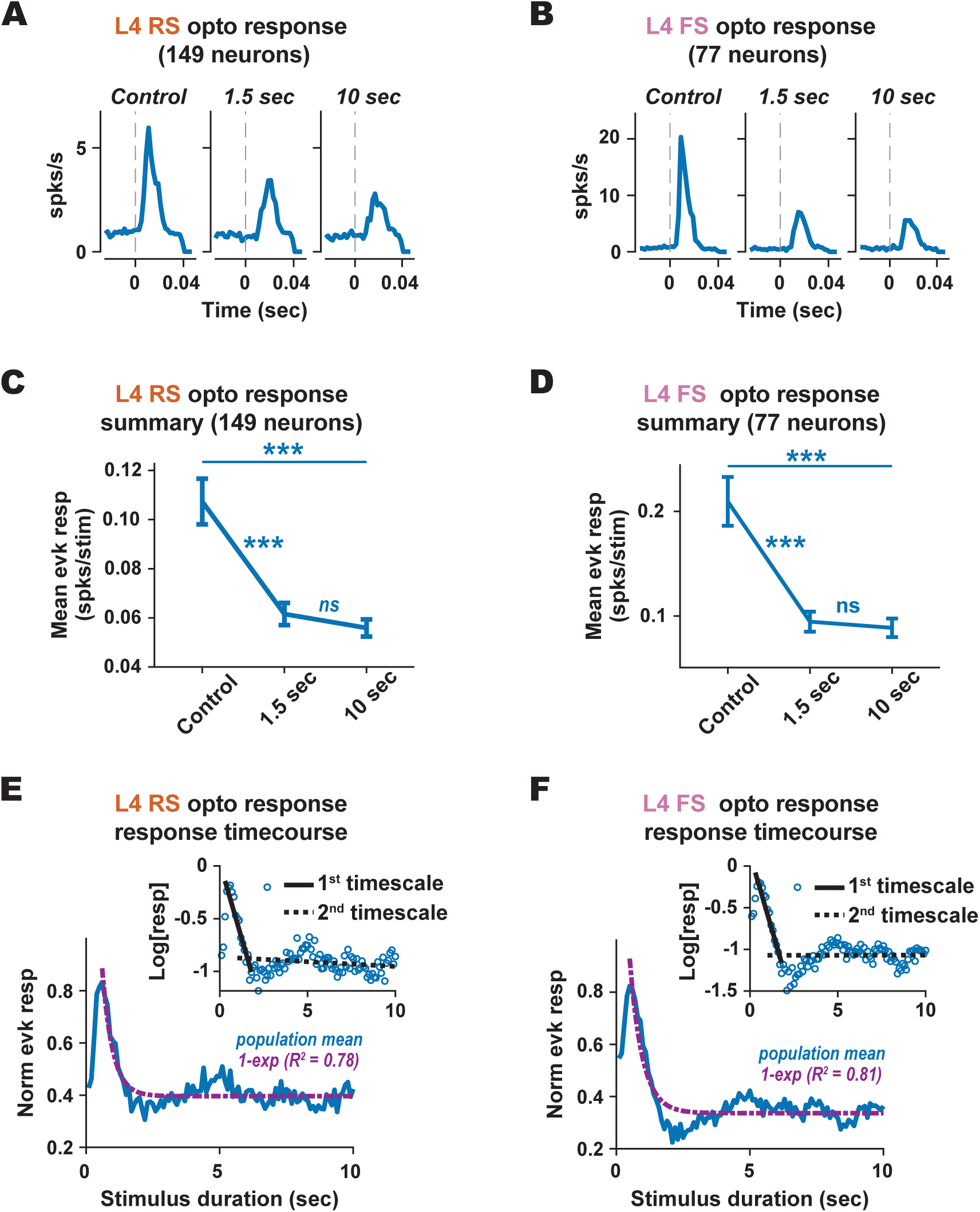
Optogenetic drive of VPm results in a single timecourse of adaptation in S1. **A**. Grand-averaged PSTHs of L4 RS neurons in response to 10 Hz repetitive optogenetic stimulation of VPm at the three timepoints of interest. **B.** Same as A but for L4 FS neurons. **C.** Mean ± sem evoked response of the L4 RS population to repetitive optogenetic stimuli of VPm at the three time points of interest (Control: 1.07e-01 ± 9.30e-03, 1.5 sec adapted: 6.16e-02 ± 4.52e-03, 10 sec adapted: 5.59e-02 ± 3.49e-03. Control versus 1.5 sec adapted: p = 3.89e-13; 1.5 sec adapted versus 10 sec adapted: p = 2.63e-01, Control versus 10 sec adapted: p = 1.76e-15, all statistical comparisons were performed using the Wilcoxon signed-rank test with a Bonferroni correction; *** indicates p < 3.3e-04). **D.** Same as C, but for the L4 FS population (Control: 2.09e-01 ± 2.33e-02, 1.5 sec adapted: 9.46e-02 ± 9.58e-03, 10 sec adapted: 8.87e-02 ± 8.84e-03. Control versus 1.5 sec adapted: p = 3.64e-09; 1.5 sec adapted versus 10 sec adapted: p = 3.99e-01, Control versus 10 sec adapted: p = 2.87e-11, all statistical comparisons were performed using the Wilcoxon signed-rank test with a Bonferroni correction; *** indicates p < 3.3e-04). **E.** Population-averaged evoked response for the L4 RS population over the timecourse of repetitive optogenetic stimulation of VPm overlaid with a single exponential model fit. Inset: semi-log plot of the normalized evoked response; the black lines denote the linear fit over one timescale plus a constant, suggesting single exponential behavior (τ_1_ = 0.39 sec, constant = 0.40). **F.** Same as E, but for the L4 FS population (τ_1_ = 0.47 sec, constant = 0.34).

Through these control experiments, we find that optogenetic drive of VPm that eliminates the longer timescale adaptation in both firing rate and synchrony in the VPm neuron population also eliminates the longer timescale adaptation in S1 layer 4 neurons. This suggests that the previous observations of the longer timescale adaptation in S1 neurons during sensory drive are inherited from VPm thalamus, consistent with a growing notion of VPm as the source of much of the adaptation observed in S1 (Wright et al., 2021).

### Adaptation over multiple timescales degrades stimulus detection for an ideal observer of cortical activity

Having explored thalamocortical activity across multiple timescales of sensory adaptation, we next sought to investigate the functional consequences of these timescales. In previous studies of the vibrissa pathway, we have shown that repetitive stimulation lasting as brief as 1-2 seconds is sufficient to not only reduce the overall responsiveness of the cortical population but shift the neuronal responses towards an adaptive trade-off that favors sensory discrimination over detection (Wang et al., 2010b; Ollerenshaw et al., 2014). To determine if this holds true for longer timescales of adaptation, we probed cortical responses either in the presence or absence of a preceding adapting stimulus and leveraged signal detection theory to ascertain the ability of an ideal observer to detect a signal (stimulus present) versus noise (stimulus absent; Figure 8A). Although the results in Figure 2 regarding the decrease in the magnitude of the sensory evoked cortical response would suggest a corresponding decrease in detectability, this is ultimately determined by both the magnitude of the response and the corresponding variability, relative to ongoing cortical activity. To directly test this, the cortical response to a 300 deg/s punctate stimulus and the corresponding detectability were evaluated in isolation (Control) and in the presence of preceding 10 Hz adapting stimuli of 1.5 second or 10 second durations.

**Figure 8.**
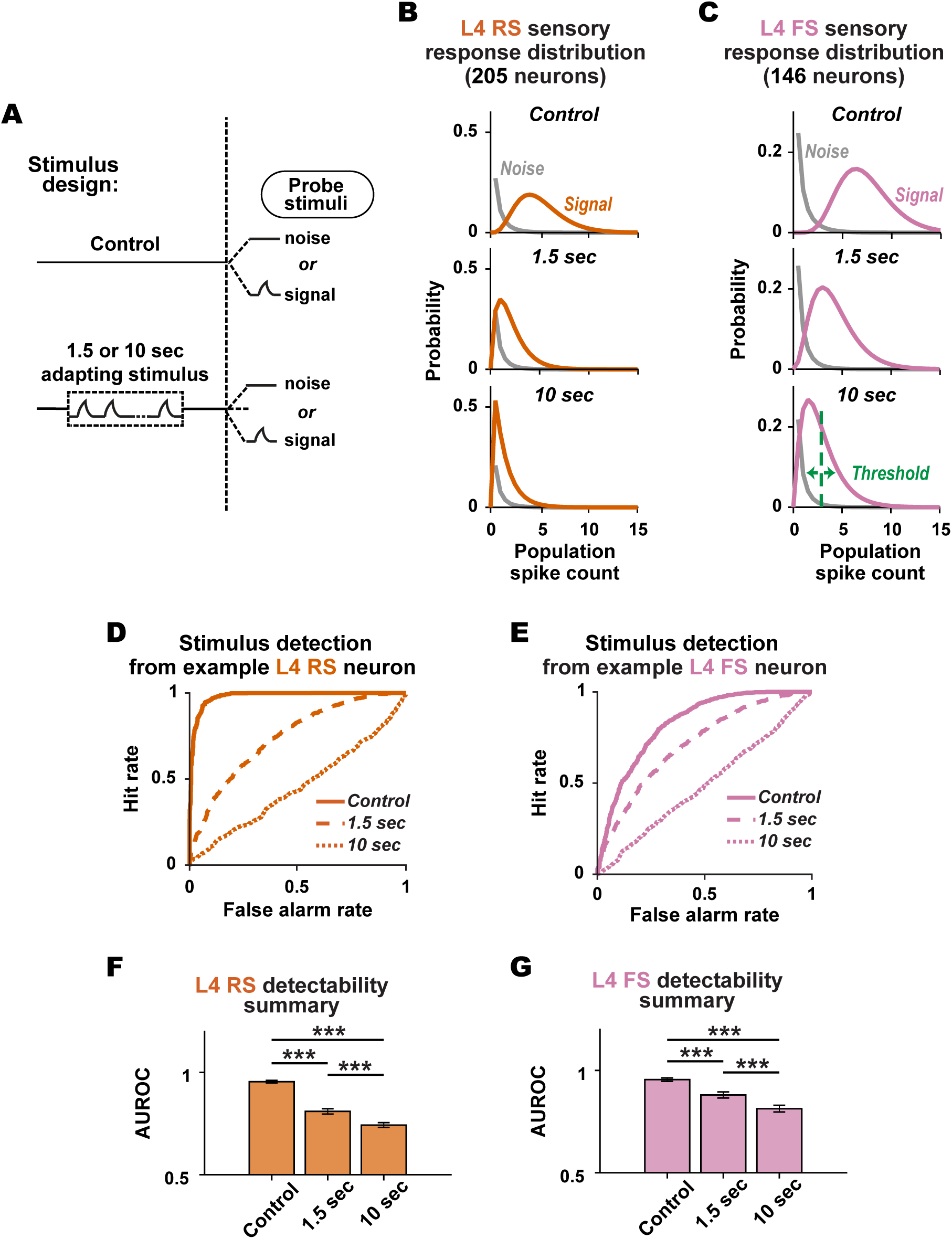
Adaptation degrades stimulus detection for an ideal observer of cortical activity. **A**. Cortical responses to sensory inputs of 300 deg/s were probed either in the presence or absence of a preceding adapting stimulus. **B**. Parametric signal (stimulus present) and noise (stimulus absent) population spike probability distributions for L4 RS neurons when the probe stimulus is preceded by the Control (top), 1.5 sec adapted (middle), or 10 sec adapted (bottom) stimulus. Parametric fits were generated using a Gamma distribution on the L4 RS population response (see Methods). **C**. Same as in B, but for the L4 FS neuron population. **D**. Receiver operator characteristic (ROC) curves of the sensory response of an example L4 RS neuron when the stimulus is preceded by the Control, 1.5 sec adapting, or 10 sec adapting stimuli. **E**. Same as D, but for the L4 FS population. **F**. Mean ± sem performance of an ideal observer of L4 RS activity in detecting the signal versus noise in conditions preceded by Control, 1.5 sec adapting, and 10 sec adapting stimuli. The performance is quantified by the area under the receiver operator characteristic curve (AUROC) performed across neurons; AUROC for control: 0.96 ± 6.36e-03, 1.5 sec adapted: 0.81 ± 1.31e-02, 10 sec adapted: 0.74 ± 1.18e-02; Control versus 1.5 sec adapted: p = 5.05e-23, 1.5 sec versus 10 sec: p = 5.31e-07, Control versus 10 sec adapted: p = 5.68e-29; all statistical comparisons were performed using the Wilcoxon signed-rank test with a Bonferroni correction; *** indicates p < 3.3e-04). **G**. Same as F but for L4 FS neurons (AUROC for control: 0.96 ± 8.04e-03, 1.5 sec adapted: 0.88 ± 1.45e-02, 10 sec adapted: 0.81 ± 1.63e-02; Control versus 1.5 sec adapted: p = 4.76e-10, 1.5 sec versus 10 sec: p = 4.76e-09, Control versus 10 sec adapted: p = 1.37e-17), all statistical comparisons were performed using the Wilcoxon signed-rank test with a Bonferroni correction; *** indicates p < 3.3e-04).

To quantify cortical sensory responses under the signal detection framework, we constructed parameterized probability distributions of the signal and noise spike counts resulting from the probe stimulus preceded by the Control and adapting stimulus conditions. The signal distributions were estimated by drawing samples from gamma distribution fits of the observed cortical spike counts within a 30 ms post-stimulus response window for a 300 deg/s stimulus. The noise distributions were estimated from baseline activity 30 ms prior to stimulus onset (see Methods). We find that adaptation pushed the signal distribution of the evoked response closer to the noise distribution at each time point of interest for both L4 RS and FS populations (Figures 8B & 8C, respectively).

To predict the potential impact of adaptation on stimulus detectability, we quantified the extent to which an ideal observer of cortical spiking activity accurately detects the presence of a signal from background noise when the signal is preceded by the Control and adapted conditions. Thus, we applied a sliding threshold on the parametric distributions to generate a Receiver Operator Characteristics (ROC) curve which shows the hit rate vs the false alarm rate for an ideal observer of the neuronal activity. For ROC curves, a perfect diagonal indicates that the ideal observer is operating at chance, and deviations above the diagonal indicate improved performance. Figures 8D & 8E show the ROC curves for an example L4 RS and FS neuron, respectively, for each of the three stimulus conditions. For both populations, adapting stimuli preceding the probe stimulus brought the ROC curve closer to the diagonal, reflective of a loss in theoretical detection performance. This loss was greatest when the probe stimulus was preceded by 10 seconds of an adapting stimulus.

To quantify this loss in theoretical detection performance, we measured the area under the ROC curve (AUROC) for each neuron. An AUROC value of 1 corresponds to zero overlap between signal and noise distribution (an ideal detector). An AUROC value of 0.5 corresponds to complete overlap between distributions, hence the ideal observer operating at chance. We found that the detectability of both RS and FS neurons significantly decreased when preceded by Control versus 1.5 second adapting stimuli, and further decreased when preceded by 10 second adapting stimuli (Figures 8F & 8G). Consistent with the observation in Figure 1F that RS neurons had lower adaptation ratios than FS neurons, we found that the theoretical detectability was generally lower for RS neurons as compared to FS neurons.

### Adaptation over multiple timescales enhances stimulus discrimination for an ideal observer of cortical activity

We next evaluated the functional consequences of adaptation on stimulus discrimination across multiple timescales by presenting probe stimuli of varying velocities (50, 300, 450, 600, or 1200 deg/s), preceded by either Control, 1.5 second, or 10 second adapting stimuli. First, in terms of the impact of adaptation on the velocity sensitivity properties in the cortex, adaptation attenuated the response magnitude across all velocities for cortical populations, although differently for different velocities. The velocity sensitivity was further reduced for both the RS and FS populations when the probe stimuli were preceded by the 10-sec10 second adapted stimulus compared to the 1.5 second adapted stimulus (Figures 9A and 9B).

**Figure 9.**
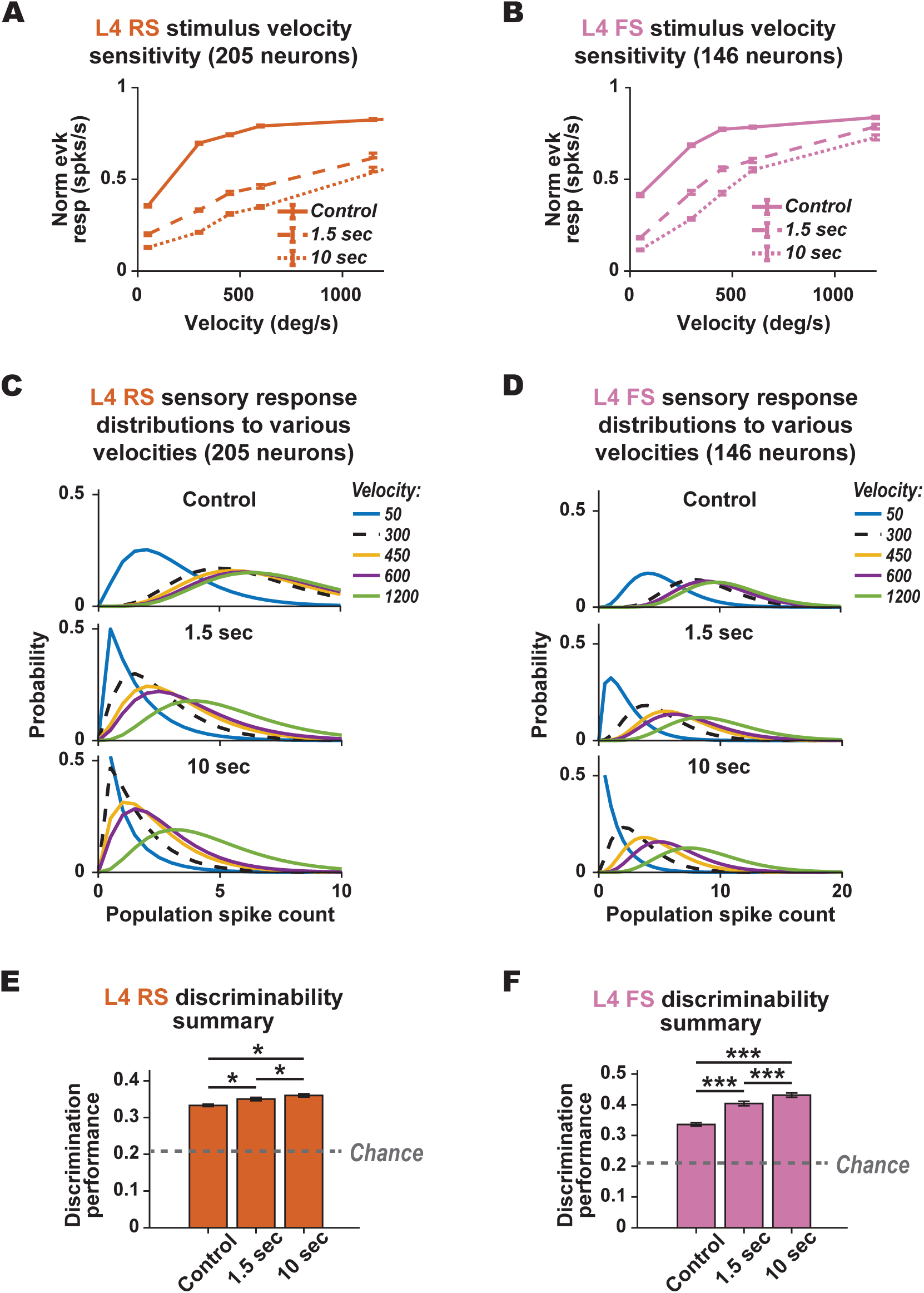
Adaptation enhances stimulus discrimination for an ideal observer of cortical activity. **A**. Stimulus response curves of the L4 RS population response to probe stimuli of varying velocities preceded by Control, 1.5 sec adapting, or 10 sec adapting stimuli. **B.** Same as A, but for the L4 FS population. **C.** Parametric distribution of L4 RS sensory responses to probe stimuli of varying velocities when the probe stimulus is preceded by the Control (top), 1.5 sec adapting (middle), or 10 sec adapting (bottom) stimuli. Parametric fits are generated using a Gamma distribution of the RS population response (see Methods). **D**. Same as C but for the L4 FS population. **E**. The performance of an ideal observer of L4 RS activity in discriminating between sensory stimuli of varying velocities in conditions preceded by Control, 1.5 sec adapting, and 10 sec adapting stimuli (mean +/- 95% confidence intervals from re-sampling neurons with replacement; see Methods). The performance is quantified by the accuracy of an LDA classifier in discriminating between all velocities (Control: 0.33 ± 3.1e-03, 1.5 sec adapted: 0.35 ± 4.7e-03, 10 sec adapted: 0.36 ± 4.3e-03. Control versus 1.5 sec adapted: p = 1.7e-02, 1.5 sec versus 10 sec, p = 1.7e-02, Control versus 10 sec adapted: p = 1.7e-02; all statistical comparisons of confidence intervals were performed with a Bonferroni correction). **F**. Same as E but for the FS population (mean +/- 99.9% confidence intervals from re-sampling neurons with replacement; Control: 0.34 ± 5.7e-03, 1.5 sec adapted: 0.41 ± 7.3e-03, 10 sec adapted: 0.43 ± 6.9e-03; Control versus 1.5 sec adapted: p = 3.3e-04, 1.5 sec versus 10 sec, p = 3.3e-04, Control versus 10 sec adapted: p = 3.3e-04).

Taking the perspective of an ideal observer, we examined the extent to which adaptation over multiple timescales affects discrimination between whisker deflections at different velocities. Shown in Figures 9C and 9D are parametric fits of the experimentally observed spike count distributions for the RS and FS populations, respectively. Qualitatively, the populations’ spike counts were relatively high in the Control conditions, but with highly overlapped and unordered distributions. However, while increasing pre-stimulus adaptation attenuated the population spikes counts, the distributions became more separated and ordered (Figures 9C & 9D).

To quantify this effect, we compared the performance of the ideal observer in discriminating between different deflection velocities when preceded by Control, 1.5 second and 10 second adapting stimuli. For each stimulus condition, we randomly selected 25 neurons from each L4 RS and L4 FS population and combined stimulus trials to form sensory response distributions of the cortical responses to various velocities (parameterized gamma distributions, see Methods). To assess discrimination performance, we implemented a holdout validation approach, reserving data from 20% of trials for testing while using the remaining 80% of trials for training. Linear discriminant analysis (LDA) classifiers were then applied to infer the most likely velocity presented based on the observed cortical response, and discrimination performance was evaluated by the accuracy of the ideal observer in making correct stimulus classifications (see Methods). This process was iterated 1000 times to comprehensively assess discrimination performance for each stimulus condition (Control, 1.5 second, and 10 second adapted) across various combinations of neurons and trials. Given that all five velocities were presented with equal probability, a value of 0.2 reflects the ideal observer operating at chance. We found that increasing adaptation significantly improved overall discrimination performance in both RS and FS populations (Figures 9E & 9F), across the early and later stages of the adaptation. Taken together, our results suggest that adaptation over multiple timescales continues to facilitate better discrimination performance by reducing the overlap between stimulus distributions by trading off the overall sensitivity to repetitive stimulation.

## DISCUSSION

In this study, we discovered two clear and distinct timescales of rapid adaptation across the thalamocortical circuit over the course of seconds. Several other studies have examined the properties of adaptation over a range of timepoints using various durations of repetitive stimulation (Dragoi et al., 2000; Patterson et al., 2013; Latimer et al., 2019), white noise stimulation, (Fairhall et al., 2000; Lundstrom et al., 2010) and modeling approaches (Latimer and Fairhall, 2020). Here, the ability to record simultaneously from the thalamic VPm population as well as both the cortical L4 RS and FS populations allowed us to precisely examine transformations of adaptation across the thalamocortical junction through the spiking response amplitudes, thalamic population synchrony, and synaptic efficacy.

Numerous studies have identified potential circuit mechanisms that could underlie the differential adaptation from thalamus to cortex. Potential drivers of the observed rapid cortical sensory adaptation include depression of thalamocortical (TC) synapses (Chung et al., 2002) and/or synaptic depression/facilitation of excitatory/inhibitory intracortical (IC) projections (Beierlein and Connors, 2002; Reig et al., 2006), and neuronal population mechanisms such as thalamic firing rate and population synchrony, all of which may collectively shape the IC balance of excitation and inhibition. One prominent view is that short-term depression of thalamocortical synapses plays a dominant role in this type of cortical adaptation (Chung et al., 2002; Castro-Alamancos, 2004b; Katz et al., 2006). However, the rapid synaptic depression of the TC synapse reported in Chung et al., 2002 does not rule out the contribution of thalamic population synchrony observed here in shaping S1 adaptation, which was not measured in the relevant studies. Further, subsequent studies suggest that thalamocortical desynchronization plays perhaps an even more prominent role than short-term changes in the TC or IC synapses (Wright et al., 2021). In fact, many lines of evidence have demonstrated the profound effect that thalamic spike timing and synchrony has on S1 processing either under anesthesia (Pinto et al., 2000; Wang et al., 2010b; Whitmire and Stanley, 2016) or wakefulness (Wright et al., 2021; Borden et al., 2022). The optogenetic control experiments here do not rule out the possibility of TC synaptic depression playing a role in the fast timescale adaptation, but previous findings (Wright et al., 2021) suggest that this role may be overshadowed by the adaptive reductions in thalamic firing rate and synchrony that are also clearly present in the adaptation at fast timescales (0-1.5 seconds). What the optogenetic control experiments here do show is that the monotonic slow desynchronization of the thalamic population beyond the first few seconds (2-10 seconds) seems to play a dominant role in modulating cortical adaptation at longer timescales, with little or no influence from TC synaptic depression on this timescale. These results together suggest the possibility of a complex interaction between distinct mechanisms, and that multiple mechanisms are likely co-activated over distinct periods of time to contribute to the cellular dynamics that we observed.

Differential adaptation of the putative excitatory and inhibitory cortical neurons observed here likely shapes overall cortical excitability and the integration properties of ascending sensory signals from thalamus. Importantly, this effect has been shown during wakefulness, but the extent of the differential adaptation is strongly shaped by the nature of the adapting stimulus itself (Wright et al., 2021). The more profound depression of intracortical inhibitory synapses as compared to excitatory synapses during persistent stimulation has been shown to skew the E-I balance towards excitation (Gabernet et al., 2005; Heiss et al., 2008). This reduction in feedforward inhibition was shown to expand the cortical window of integration and affect how inputs are summed across multiple timescales of adaptation. This mechanism could potentially regulate the effects of the thalamic desynchronization observed here, offsetting the loss of thalamic synchrony through a loss in cortical sensitivity to thalamic synchrony. However, in previous work, we have shown that the desynchronization observed in thalamus during adaptation outweighs the changes in cortical integration (Wang et al., 2010b).

This study highlights the important contributions of the rapid adaptation of thalamus in shaping cortical adaptation. What is the origin of the adaptative properties observed in sensory thalamus? For the fastest timescale component of the VPm adaptation, given the high firing rates of neurons in the brainstem (Ahissar et al., 2000), it is likely that depression of the synaptic inputs from brainstem to thalamus play an important role in the rapid decrease in thalamic firing rate, as well as the desynchronization in the thalamic population at this timescale. One important finding here was that the longer timescale adaptation observed in S1 appears to be largely inherited from VPm. When VPm thalamus was optogenetically driven, this eliminated the slow decrease in thalamic firing rate and synchronization (Figure 6), which subsequently eliminated the slower, later phase of adaptation in S1 (Figure 7). One distinct possibility is that the direct optogenetic drive of VPm overcomes intrinsic properties of the thalamic neurons that are responsible for the slow drift. In brain slice experiments, thalamic neurons have been shown to exhibit a slow after-hyperpolarization that persists for several seconds after controlled depolarization (Goaillard and Vincent, 2002). The light activation of Channelrhodopsin in the thalamic neurons likely overcomes these intrinsic cell properties, eliminating the slow after-hyperpolarization and the corresponding decrease in firing rate and synchronization of the thalamic population. However, further experiments are needed to more fully understand this process in-vivo and to better understand the interaction between intrinsic properties of thalamic neurons and the ascending input from the brainstem.

The adaptation in firing rate across VPm and S1 evolved on similar timescales as reflected qualitatively in Figure 1E and quantitatively through comparable time constants, although the time constant associated with the slower component was longer in S1 as compared to VPm. However, the extent of the adaptation was more extreme in S1 as compared to VPm, as reflected in the steady-state firing rate relative to the non-adapted firing rate at the onset of the adaptation, consistent with previous observations (Chung et al., 2002; Khatri et al., 2004; Gabernet et al., 2005). Further, within S1 the RS neurons adapted more than the FS neurons, also consistent with previous reports (Wright et al., 2021). It is also the case that despite having a similar qualitative nature, the slower timescale component of the thalamic synchrony has a significantly longer associated time constant as compared to the cortical or thalamic firing rates. This suggests that the thalamic synchrony continues to diminish even when thalamic and cortical firing rates have bottomed out. However, given the nonlinear sensitivity of cortex to the synchrony of the thalamic input (Bruno and Sakmann, 2006; Wang et al., 2010b; Wright et al., 2021), even the small decrease in thalamic synchrony observed on the longer timescale here does not preclude thalamic synchrony from being a primary role player in the corresponding cortical firing rate adaptation.

In this study, we were able to capture multiple timescales of adaptation using a 10 Hz repetitive stimulation over 10 seconds. It is important to note, however, that adaptation is a change in function that takes time to develop and also dissipate. It has been shown that the temporal scales over which adaptive dynamics build up and recover are distinct, and also highly dependent upon the duration of exposure to repetitive stimulation (Dragoi et al., 2000; Chung et al., 2002; Patterson et al., 2013). In general, brief exposure produces adaptation that develops and recovers rapidly (Bonds, 1991; Nelson, 1991; Müller et al., 1999), while more prolonged stimulation can result in slower and more lasting forms of changes (Greenlee et al., 1991). Additionally, thalamic and cortical neurons also exhibit diverse timescales of responsiveness to repeated stimulation (Chung et al., 2002). Therefore, as a rule of thumb, an inter-trial interval of at least 4 times the duration of adapting stimulus train (10 seconds) was utilized in this study, while acquiring at least 100 trials of repeated measurements of the adaptation dynamics. As a control, we also assessed the difference between the magnitude of the response to the first deflection of the adapting train at the beginning and end of a stimulus block. We did not observe a significant difference in this measure, suggesting that the majority of the neurons returned to the baseline condition prior to the initiation of the next trial, although longitudinal effects cannot be completely ruled out. For a subset of the data, we also probed the recovery for each thalamic and cortical neuron and found that most of the cells showed a full recovery to the baseline condition. Finally, it is important to note that different aspects of the adapting stimulus itself, such as frequency content and velocity, can strongly shape the nature of the adaptation as we have previously shown (Zheng et al., 2015). Although the exact timescales of the adaptation likely vary with adapting stimulus properties, we believe the primary finding of multiple distinct timescales of adaptation to be general.

The direct assessment of the transformation of adaptation across the thalamocortical junction in this study is facilitated by our simultaneous multi-channel electrode recordings of putatively connected pairs of thalamocortical neurons. From the total number of topographically aligned thalamocortical pairs recorded in this study, we identified approximately 33% as putatively connected, similar to but slightly lower than that previously described in the cat visual pathway (42%, Reid and Alonso, 1995). This difference in inferred connectivity can likely be attributed to the conservative nature of our inference methodology, which was explicitly designed to tolerate misses over false alarms (Liew et al., 2021). Further, the inference of connectivity is very sensitive to the number of spikes used in the analysis; the nature of the multi-electrode arrays used in this study resulted in sampling of neurons with highly diverse firing rates, including those with relatively low firing rates that precluded the inference of connectivity. These two factors likely resulted in excluding pairs that were actually connected from the analyses here. When parsed based on S1 cell-type classification, we further found a higher probability of likely connectivity among VPm-FS (46.8%) as compared to VPm-RS (24.8%) pairs, consistent with previous findings (Bruno and Simons, 2002; Bruno and Sakmann, 2006).

Given that this study was performed in the somatosensory pathway of isoflurane-anesthetized mice, it is important to acknowledge some of the limitations in the findings. The large majority of in-vivo studies of the relative role of short-term depression of thalamocortical and intracortical synapses have been conducted under anesthesia (Heiss et al., 2008; Chung et al., 2019), and the current study cannot rule out the potential influence of TC synaptic depression in the early, fast timescale component of adaptation. In contrast, the adaptive changes in synchronous thalamic firing have recently been reported to play a more prominent role than synaptic depression in the TC and IC synapse under wakefulness (Wright et al., 2021). One possibility is that the overlap between distinct mechanisms during wakefulness is smaller compared to the condition under anesthesia. It was reported that the thalamocortical circuit may be in an elevated baseline firing state during wakefulness (Castro-Alamancos, 2004a), hence the suggestion that TC synapses might already be depressed in this state. When TC synaptic depression is fully engaged during the high thalamic firing rates associated with wakefulness, thalamic desynchronization may dominate and exert an even greater effect on regulating the nature of information conveyed downstream to cortical targets. However, to fully disentangle these effects and the relative contribution from each mechanism, further techniques would need to be employed, such as selective optogenetic manipulation of TC and CC synaptic depression combined with simultaneous, multi-site recordings, beyond the scope of the current study.

Finally, the results here suggest that although multiple mechanisms may interact and transition dynamically at various timescales, adaptation over the timecourse that we examined in this study improve theoretical discrimination at the expense of detection. This similarity in function supports the possibility of a unifying framework across disparate mechanisms and timescales, which is to allow the brain to maintain a flexible representation in response to stimulus statistics that span a wide range in our natural environment. Moreover, these results further support Barlow’s efficient coding hypothesis such that the reduction in responsiveness of neuronal response in the TC circuit is not a negative attribute that signifies a loss of sensitivity to stimuli, but instead a mechanism that could mediate an adaptive trade-off in coding strategy to put the brain, in this case the primary sensory cortex, in a state that optimizes neural coding efficiency (Barlow, 1961).

## Acknowledgements

This work was supported by NIH National Institute of Neurological Disorders and Stroke (NINDS) BRAIN Grant R01NS104928 and NINDS BRAIN Grant RF1NS128896. YJL was supported by a Georgia Tech-Emory-PKU Global Biomedical Engineering Fellowship. EDD was supported by a National Science Foundation Graduate Research Fellowship and the Howard Hughes Medical Institute through the James H. Gilliam Fellowships for Advanced Study program. YZ was supported by STI2030-Major Projects 2022ZD0204900. The authors thank Eunji Cheong and Cheong lab members at Yonsei University for helpful conversations at various stages of this work.

